# From duplication to divergence: single-cell insights into transcriptional and *cis*-regulatory landscapes

**DOI:** 10.1101/2025.05.07.652760

**Authors:** Xiang Li, Xuan Zhang, Robert J. Schmitz

**Author notes:** The author(s) responsible for distribution of materials integral to the findings presented in this article in accordance with the policy described in the Instructions for Authors (https://academic.oup.com/plcell/pages/General-Instructions) is Xiang Li.

## Abstract

Gene duplication is a major source of evolutionary innovation, enabling the emergence of novel expression patterns and functions. Leveraging single-cell genomics, we investigated the transcriptional and *cis*-regulatory landscapes of duplicated genes in cultivated soybean (*Glycine max*), that has undergone two rounds of whole-genome duplication. Our analysis revealed extensive diversity of transcriptional profiles within and across tissues among duplicated gene pairs. Within-tissue divergence was largely attributable to genetic variation in their associated accessible chromatin regions (ACRs), where *cis*-regulatory elements reside, whereas cross-tissue divergence was more likely shaped by dynamics in ACR chromatin accessibility profiles across tissues. Distinct duplication mechanisms also likely give rise to different types of *cis-*regulatory variants, contributing variably to transcriptional divergence. By comparing ACRs associated with gene sets derived from the two rounds of whole-genome duplications and sharing a common ancestral gene, we found that most ACRs retained one or multiple corresponding duplicated sequences in which mutations gradually accumulated over time, while a subset likely arose *de novo*. Lastly, we traced the evolution of cell-type-specific expression and cell-type-specific ACRs within duplicated gene sets, illustrating a powerful framework for identifying candidate regulatory regions driving cell-type-specific expression. Collectively, our findings highlight the important role of *cis-* regulatory evolution in shaping transcriptional divergence in a spatiotemporal manner, uncovered with the resolution of single-cell genomics.

## Introduction

Gene duplication is a crucial driver of evolution, providing the raw materials for new gene functions and expression patterns (Lynch and Conery 2000; Birchler and Yang 2022). It arises through small-scale duplication, such as tandem or proximal duplications, or through whole-genome duplication that replicates entire genomic segments simultaneously (Zhang 2003; Kuzmin et al. 2022). These mechanisms generate duplicated genes with varying evolutionary rates, essentiality (e.g. whether a gene is required for survival or normal growth), and functions (Freeling 2009; Qiao et al. 2019; Shi et al. 2020; Cai and Des Marais 2024; Benoit et al. 2025). In plants, a large proportion of annotated genes have duplicates, mostly derived from whole-genome duplication, consistent with the prevalence of palaeopolyploidy in land plants (Wendel 2000; Blanc and Wolfe 2004; Panchy et al. 2016; Tiley et al. 2016). The retention of these duplicates has been instrumental in shaping plant genomes and contributing to important agricultural traits (Panchy et al. 2016; Salman-Minkov et al. 2016; Benoit et al. 2025).

Retained duplicates may be subject to relaxed selective constraints, with one or both copies sometimes acquiring new functions (neofunctionalization) that contribute to adaptation or partitioning ancestral functions (subfunctionalization). However, functional divergence is not always necessary for their retention and typically unfolds over extended evolutionary timescales (Panchy et al. 2016; Birchler and Yang 2022). Multiple mechanisms have been proposed to account for the dispensability of retained duplicates (Innan and Kondrashov 2010; Panchy et al. 2016; Qiu et al. 2020). Moreover, factors such as selection and genetic drift can also influence gene retention (Panchy et al. 2016; Iohannes and Jackson 2023). On the other hand, this redundancy invariably complicates genotype-phenotype associations and limits the transferability of genetic findings across closely related species (Iohannes and Jackson 2023; Benoit et al. 2025). Despite their evolutionary and functional significance, the dynamics of gene duplication and diversification over short evolutionary timescales remain insufficiently explored (Benoit et al. 2025).

Divergent expression between duplicated genes is a widespread phenomenon and provides crucial insight into their evolution. In Arabidopsis, 70% of duplicated gene pairs exhibit significant transcriptional differences (Ganko et al. 2007). Similarly, approximately 50% of soybean duplicated genes, derived from whole-genome duplication, show transcriptional divergence (Roulin et al. 2013). This divergence is likely driven by genetic variants within regulatory regions (Wray et al. 2003; Tran et al. 2024), particularly *cis*-regulatory elements that control spatiotemporal expression patterns (Wittkopp and Kalay 2012; Schmitz et al. 2022). In cotton, divergence in *cis*-regulatory sequences between its diploid progenitors contributes to transcriptional divergence in 40% of the duplicated genes (Chaudhary et al. 2009). Similar patterns have been observed in Arabidopsis (Haberer et al. 2004), soybean (Fang et al. 2023), strawberry (Fang et al. 2024) and additional cotton studies (Han et al. 2022; Hu et al. 2024). Despite these valuable insights, an underexplored question remains: what forms of transcriptional divergence and which types of *cis*-regulatory variation are maintained in duplicated genes?

Soybean (*Glycine max*) having gone through two rounds of whole-genome duplication, approximately 59 and 13 million years ago (Schmutz et al. 2010; Koenen et al. 2021; Zhuang et al. 2022), makes it an excellent model for exploring this question. Gene pairs resulting from both whole-genome and small-scale duplications have been identified in soybean (Qiao et al. 2019). Previous studies have also explored the evolutionary fates of duplicated genes in soybean, highlighting divergence in gene expression and the role of *cis*-regulatory control (Du et al. 2012; Zhao et al. 2017; Fang et al. 2023). However, these studies primarily assessed transcriptional divergence at the tissue level, often relying on differential expression as the sole metric (Fang et al. 2023), which likely oversimplifies “divergence” and overlooks differences at the cellular level.

For instance, even when two duplicated genes exhibit similar overall expression levels within a given tissue, they may display distinct expression patterns across different cell types within that tissue. This raises a critical question: do tissue-level and magnitude-based comparisons adequately capture transcriptional divergence, and to what extent does genetic variation in *cis*-regulatory regions drive such differences? Duplicated genes, which often retained copies of their associated regulatory regions, offer a powerful model for dissecting these regulatory relationships. Revealingly, variation in these regions could also result in cell-type-specific expression divergence, offering additional insight into the regulatory basis underlying cellular specificity and informing potential applications in synthetic biology. The recent availability of single-cell assay for transposase-accessible chromatin sequencing (scATAC-seq) and single-nucleus RNA sequencing (snRNA-seq) data of different soybean tissues (Zhang et al. 2024) enables the investigation of transcriptional and *cis*-regulatory divergence at single-cell resolution in a spatiotemporal context.

In this study, we integrated single-cell expression and chromatin accessibility profiles to examine transcriptional and *cis*-regulatory dynamics of duplicated genes within and across tissues. This approach provides a more nuanced understanding of transcriptional divergence and its correlation with *cis*-regulatory sequence variation, chromatin accessibility and gene duplication mechanisms. Given that *cis*-regulatory elements are typically co-duplicated with genes during whole genome duplication, we further examined their evolutionary trajectories and contributions to expression divergence in gene sets derived from two rounds of whole-genome duplication and sharing a common ancestral gene. Our analysis revealed that most *cis*-regulatory elements retained one or multiple corresponding sequences across the duplicated genes, whereas a subset likely evolved *de novo*. Lastly, we explored the evolution of cell-type-specific expression within duplicated gene sets and the dynamics of their associated *cis*-regulatory regions, exemplifying how integrating cell-type-specificity with regulatory sequence conservation to identify candidate regulatory regions driving cell-type-specific expression. Overall, our findings underscore the power of single-cell genomics in revealing the divergence of expression and chromatin accessibility in duplicated genes, offering key insights into the evolution of *cis*-regulatory regions and gene duplication.

## Results

### Distinct within-tissue expression patterns of duplicated gene pairs uncovered by single-cell genomics

We identified 59,351 duplicated gene pairs in soybean using the DupGen_finder pipeline (https://github.com/qiao-xin/DupGen_finder), with common bean (*Phaseolus vulgaris*) as the outgroup. These pairs were classified into five categories: whole-genome duplication and four small-scale duplication, including tandem duplication (closely adjacent duplicated genes on the same chromosome), proximal duplication (duplicated genes located nearby on the same chromosome but separated by up to 10 other genes), transposed duplication (duplicated genes transposed to distant genomic positions, with the ancestral copy being either intra-or inter-species colinear), and dispersed duplication (duplicated genes scattered across the genome, neither neighboring nor colinear)(**Fig. 1A** and **Supplementary Table S1**). Importantly, because a single gene can experience multiple duplication events through distinct duplication mechanisms (e.g., a gene originating from whole-genome duplication may later be tandemly duplicated), we allowed redundancy in our classification, that is, the same gene may be represented in more than one duplication categories across different gene pairs. Among all identified gene pairs, whole-genome duplication derived gene pairs represented the largest proportion (50.3%), followed by dispersed duplication (37.9%) (**Fig. 1B** and **Supplementary Table S2**). The remaining 12% of duplicated gene pairs arose from tandem, proximal, and transposed duplications, with transposed duplication being the most prevalent among them (**Fig. 1B** and **Supplementary Table S2**).

**Figure 1.**
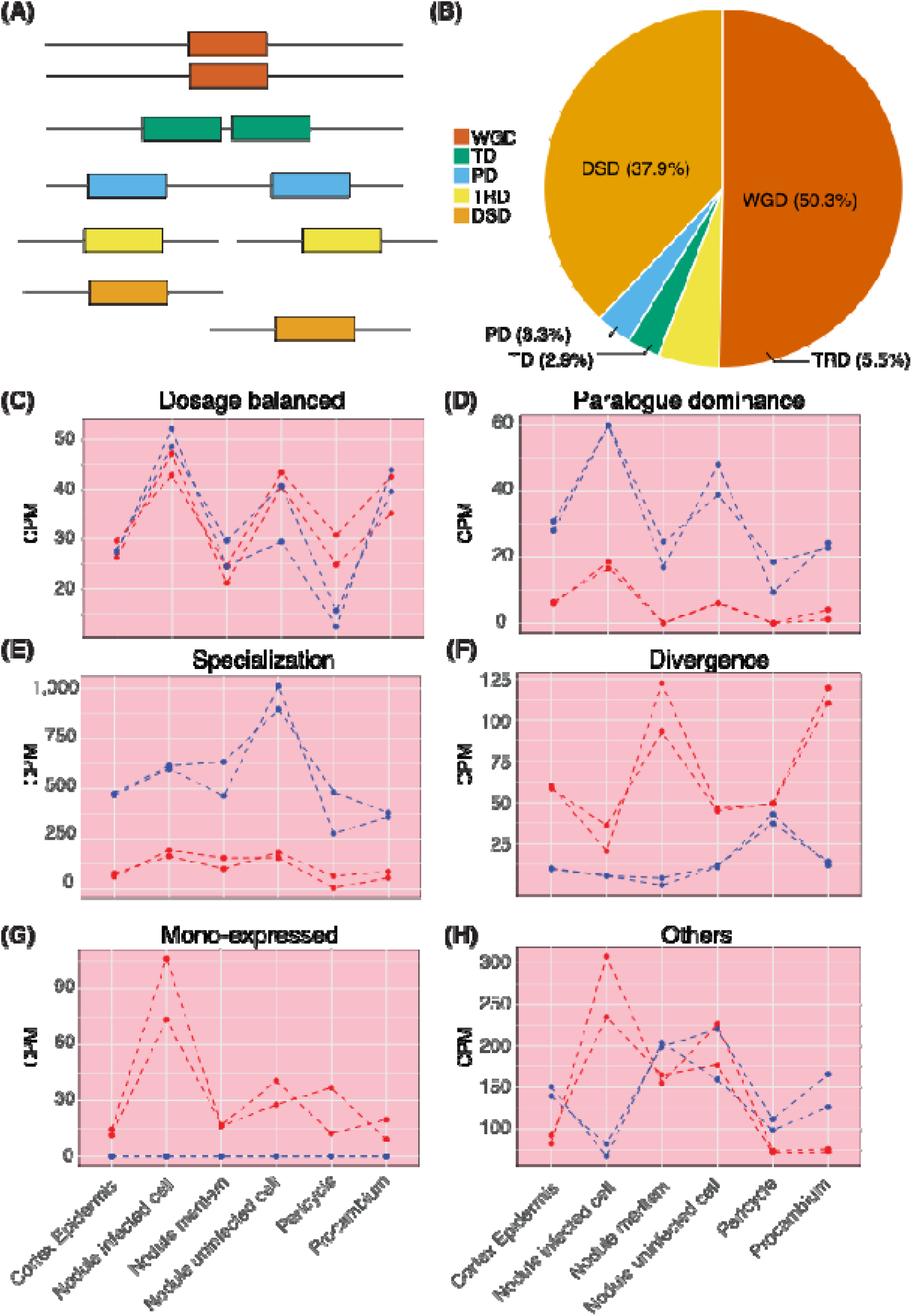
Different duplication mechanisms and distinct within-tissue expression patterns. **A)** Illustration of different gene duplication mechanisms, including whole-genome duplication (WGD), dispersed duplication (DSD), proximal duplication (PD), tandem duplication (TD), and transposed duplication (TRD). **B)** Percentages of duplicated gene pairs derived from different duplication mechanisms. **C-H)** Representative examples of duplicated gene pairs exhibiting distinct within-tissue expression patterns in early nodule tissue: (**C**) dosage balanced, (**D**) paralogue dominance, (**E**) specialization, (**F**) divergence, (**G**) mono-expressed and (**H**) others. In each panel, blue and red dots/lines represent the two genes within a duplicated pair, with two biological replicates included for each gene.

We then integrated the gene pair information with the single-cell expression profiles from each of the seven tissues - root, hypocotyl, nodule, globular stage seed, heart stage seed, cotyledon stage seed and early-maturation stage seed to examine the within-tissue expression dynamics (Zhang et al. 2024). Building on the methods developed by (Benoit et al. 2025), which measured expression correlation and differential expression of duplicated gene pairs across tissues, we adapted the approach to assess gene expression across cell types within tissue. In addition to the four previously described biologically meaningful expression patterns (Benoit et al. 2025) – (1) dosage balanced, where duplicates retain similar expression profiles and levels across cell types within a tissue (**Fig. 1C**); (2) paralogue dominance, where duplicates share similar expression profiles but exhibit consistent level of differential expression across cell types (**Fig. 1D**); (3) specialization, where duplicates display distinct expression profiles, exhibiting skewed differential expression in one or multiple cell types (**Fig. 1E**); (4) divergence, where duplicates differ in both expression profiles and levels – we introduced a fifth category (**Fig. 1F**): (5) mono-expression, in which only one gene in the pair is expressed whereas the other remains inactive in a given tissue (**Fig. 1G).** Additionally, gene pairs in which neither gene is expressed in the examined tissues are classified as non-expression, whereas those that do not fit any of the above categories are grouped as “others” (**Fig. 1H**). Notably, specialization and divergence may represent the outcomes of subfunctionalization or neofunctionalization. Among the five biologically meaningful expression patterns, the mono-expression category consistently contains the most gene pairs within each individual tissue (**Fig. 2A** and **Supplementary Table S3**). This is followed by dosage balanced, divergence, paralogue dominance, and specialization patterns, except in the hypocotyl, where divergence gene pairs outnumber dosage balanced ones, and in the early-maturation stage seed, where paralogue dominance exceeds divergence (**Fig. 2A** and **Supplementary Table S3**).

**Figure 2.**
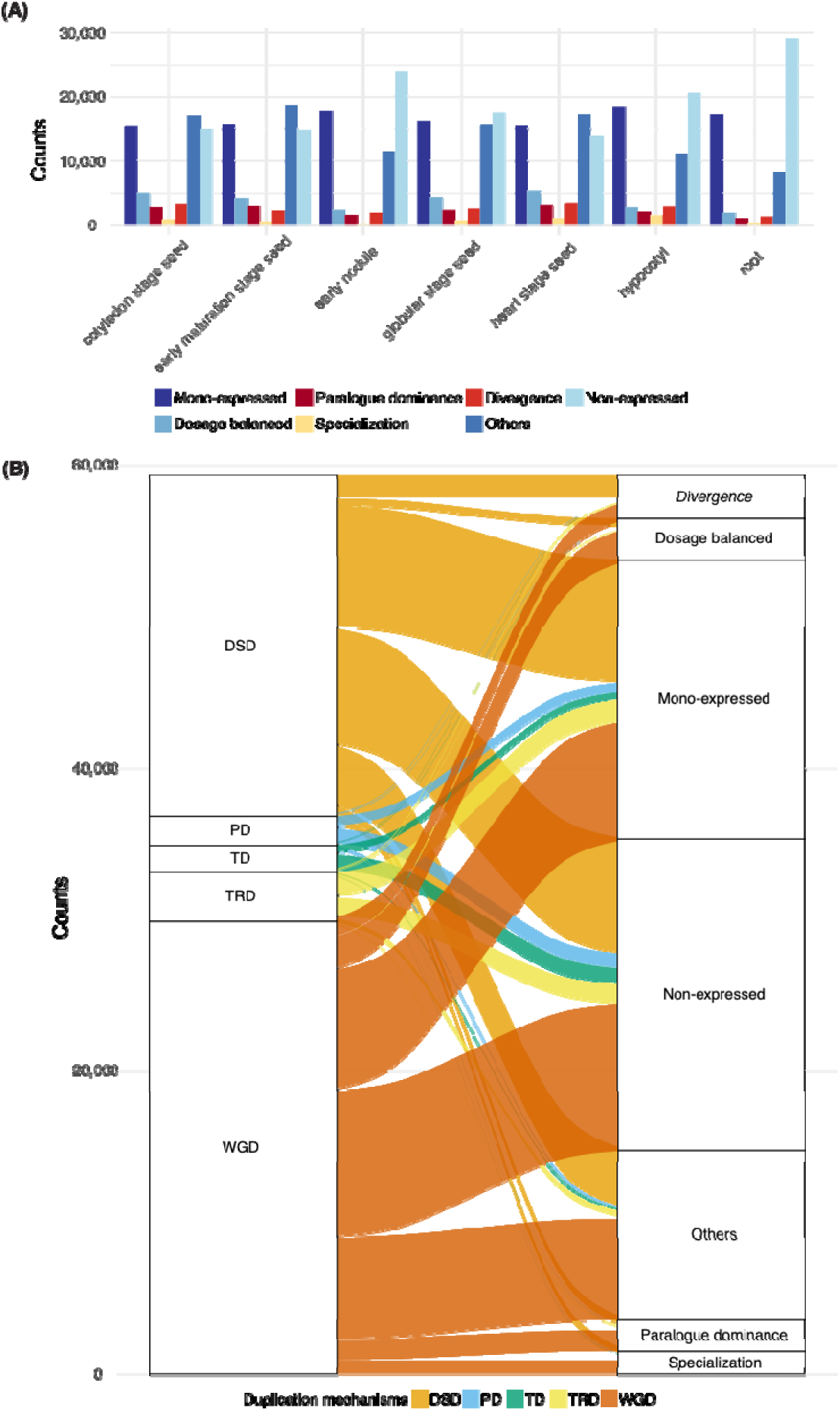
Distributions of distinct within-tissue expression patterns of duplicated gene pairs. **A)** Number of duplicated gene pairs classified by distinct expression patterns within a tissue, including mono-expressed, dosage balanced, paralogue dominance, specialization, divergence and others (as described in Fig.1), as well as non-expressed category. **B)** Relationship between gene duplication mechanisms and expression patterns. The left panel categorizes genes based on duplication mechanisms: whole-genome duplication (WGD), tandem duplication (TD), proximal duplication (PD), dispersed duplication (DSD), and transposed duplication (TRD). The right panel represents gene expression patterns. Flow width indicates the relative frequency of each transition, highlighting how different duplication mechanisms contribute to within-tissue expression variation.

When examining the expression patterns of gene pairs grouped by their duplication mechanisms, we found that those derived from whole-genome duplication and dispersed duplication exhibited similar patterns, whereas gene pairs from tandem, proximal, and transposed duplications showed greater similarity to each other (**Fig. 2B, Supplementary Fig. S1** and **Table S3**). The unpredictable, seemingly random nature of dispersed duplications (Ganko et al. 2007; Qiao et al. 2019) may explain why the expression patterns of gene pairs derived from this mechanism differ from those derived from other small-scale duplications. Additionally, the mechanisms underlying the generation of dispersed duplication remain unclear with multiple possibilities proposed (Katju and Lynch 2003; Ganko et al. 2007). Given this uncertainty, we focused on gene pairs arising from tandem, proximal, and transposed duplications, collectively referred to as regional duplications due to their tendency to remain within specific chromosomal regions, to compare their expression patterns with those of gene pairs derived from whole-genome duplication. The proportion of expressed gene pairs, where at least one gene is active in a given tissue, varied by duplication mechanisms. For instance, in early nodule tissue, about 60% of gene pairs from regional duplications (excluding transposed duplication) were not expressed, whereas only 37% of gene pairs derived from whole-genome duplication remained inactive (**Supplementary Table S3**). Among expressed regional duplicates, most exhibited mono-expression, with only a small fraction displaying other biologically meaningful expression patterns, such as dosage balanced, paralogue dominance, specialization, or divergence (**Supplementary Table S3**). These findings suggest that correlated expression patterns are more likely among whole-genome duplication derived than regional duplication derived, consistent with previous report in Arabidopsis, where duplicated genes originating from large-scale duplications and retained in duplicated segments exhibit more correlated expression patterns than those arising from small-scale duplications or those not located within duplicated segments (Casneuf et al. 2006; Qiao et al. 2019). These observations also support the gene balance hypothesis: whereas whole-genome duplication preserves the relative dosage of all interacting partners, small-scale duplication is more likely to disrupt this balance, leading to differential expression outcomes (Ohno 1970; Freeling 2009; Defoort et al. 2019; Kuzmin et al. 2022).

### Genetic variation in ACRs and its correlation with within-tissue expression patterns of duplicated gene pairs

*Cis*-regulatory variants are thought to drive transcriptional divergence in duplicated gene pairs and contribute to their retention (Iohannes and Jackson 2023); to explore this, we analyzed Accessible Chromatin Regions (ACRs), where *cis*-regulatory elements reside, from scATAC-seq data in seven tissues corresponding to the snRNA-seq data (Zhang et al. 2024). For each gene pair, we associated ACRs with duplicated genes based on their proximity.

To assess genetic variation in ACRs associated with duplicated gene pairs, we used a BLAST-based approach following the method of (Mendieta et al. 2024). For each gene pair, the ACR sequence associated with one duplicated gene was used as the query, whereas the reference was defined as the genomic region spanning from the upstream to the downstream gene of the corresponding duplicate (**Fig. 3A**). This approach accounts for the possibility that the ACR linked to the corresponding duplicate may be positioned between it and either its upstream or downstream gene. Conserved ACRs were defined based on the BLASTed ratio, calculated as the length of the BLASTed sequence within the reference region divided by the length of the query ACR. ACRs with BLASTed ratios greater than 0.1, corresponding to approximately 50 bp in length, were considered conserved (**Fig. 3B**).

**Figure 3.**
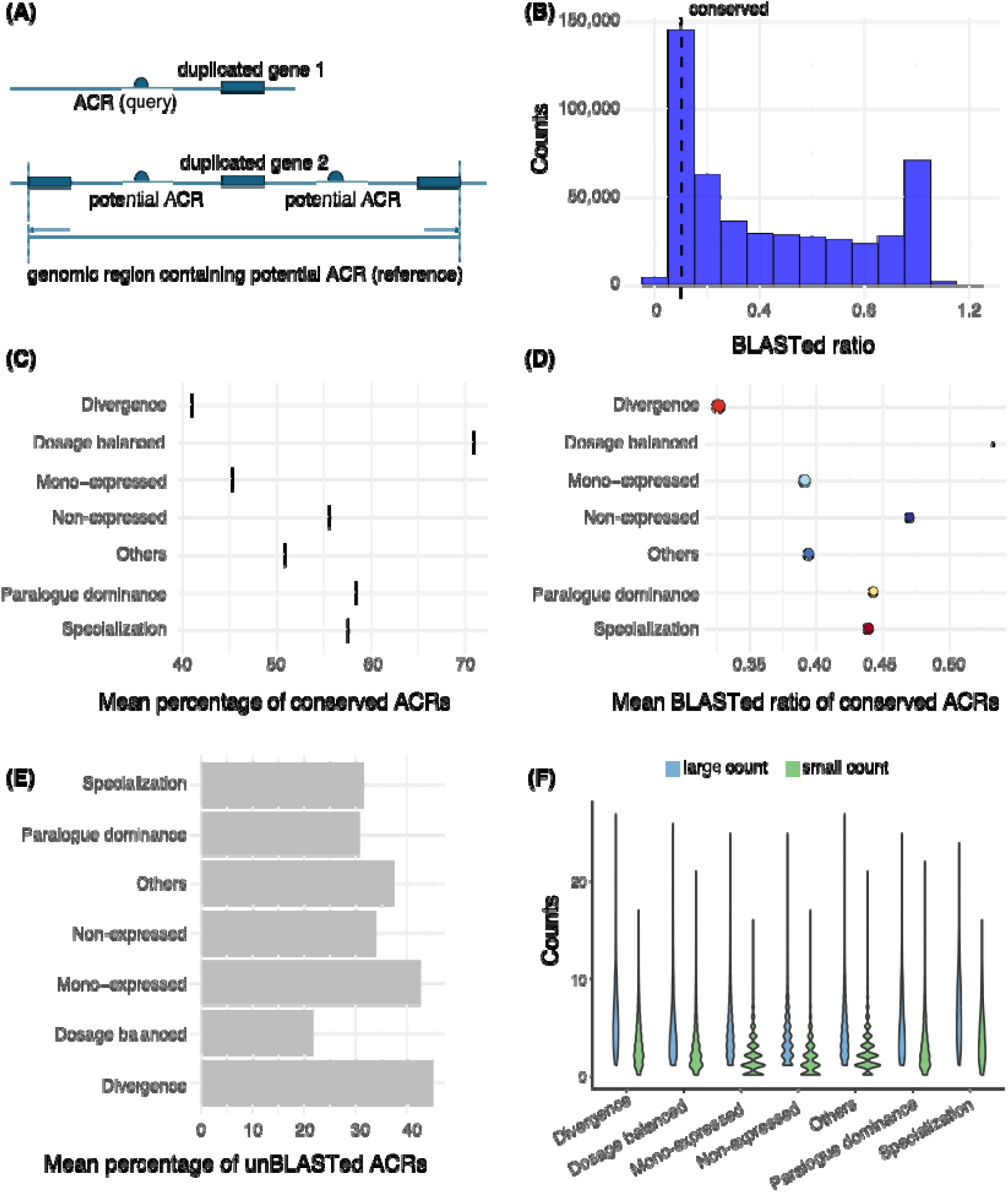
Genetic variation within ACRs associated with gene pairs with different within-tissue expression patterns. **A)** Illustration of the query and reference region for a BLAST: an ACR sequence associated with duplicated gene1 as a query, whereas the genomic region spanning from the upstream to the downstream gene of duplicated gene2 as a reference region. **B)** Number of ACRs BLASTed to the reference region in the cotyledon stage seed. ACRs with a BLASTed ratio greater than 0.1 are classified as conserved. **C)** Mean percentage of conserved ACRs (black bars) across gene pairs with different within-tissue expression patterns in the cotyledon stage seed. **D)** Mean BLASTed ratio of conserved ACRs across gene pairs with different within-tissue expression patterns in the cotyledon stage seed. The size of dots represents the mean mismatch rate of conserved ACRs across gene pairs with different within-tissue expression patterns in the cotyledon stage seed. The color of dots represents different expression patterns. **E)** Mean percentage of ACRs without a corresponding sequence in the reference region (unBLASTed ACRs) across gene pairs with different within-tissue expression patterns in the cotyledon stage seed. **F)** Number of ACRs associated with each duplicated gene within gene pairs exhibiting different within-tissue expression patterns in the cotyledon stage seed. Within each gene pair, the gene associated with a higher number of ACRs is classified into the large count group, whereas the one with fewer ACRs is classified into the small count group.

We found that gene pairs exhibiting dosage balanced had the highest percentage of conserved ACRs, whereas divergence gene pairs had the lowest (**Fig. 3C**, **Supplementary Fig. S2A** and **Table S4**). Dosage balanced gene pairs exhibit the most similar expression profiles and levels (**Fig. 1C**), whereas divergence gene pairs show great divergence (**Fig. 1F**). We also tested stricter BLASTed ratio thresholds for defining conserved ACRs (> 0.2, 0.4, 0.6 and 0.8). As expected, the proportion of conserved ACRs generally decreased with increasing thresholds (**Supplementary Table S4**). Nevertheless, the trend remained consistent: higher percentage of conserved ACR correlating with greater expression similarity between duplicated genes (**Supplementary Table S4**). Based on these observations, we focused subsequent analyses on conserved ACRs defined using the 0.1 threshold. Further examination of genetic variation between conserved ACRs and their corresponding BLASTed sequences, including BLASTed ratio and mismatch rate (calculated as the ratio of mismatches between the BLASTed sequence and the ACR query to the BLASTed sequence length), revealed greater ACR sequence divergence in gene pairs with divergence expression pattern (**Fig. 3D**, **Supplementary Fig. S2B,** and **Table S4**). We also quantified unBLASTed ACRs, those lacking detectable counterparts in their associated duplicated genes, potentially resulting from *de novo* emergence or loss of *cis*-regulatory sequences. Divergence, “others” and mono-expressed gene pairs had relatively higher proportions of unBLASTed ACRs, particularly compared to dosage-balanced gene pairs (**Fig. 3E, Supplementary Fig. S2C and Table S4**).

Another interesting observation is that, when examining the number of ACRs associated with each gene within a pair, non-expressed and mono-expressed pairs more frequently exhibited asymmetric ACR association, where only one gene had linked ACRs whereas the other had none (**Fig. 3F**, **Supplementary Fig. S2D,** and **Table S5**). In contrast, divergence gene pairs with even higher unBLASTed ratios did not exhibit this pattern (**Fig. 3F**, **Supplementary Fig. S2D,** and **Table S5**). This suggests that while ACRs might be conserved at the sequence level, their chromatin accessibility can differ between duplicates, potentially influencing whether a gene is expressed or not. Together, these observations highlight a correlation between expression divergence and ACR dynamics, shaped both by sequence divergence between conserved ACRs and their corresponding counterparts, and by variation in accessibility despite sequence conservation.

When further examining genetic variation in ACRs associated with gene pairs grouped by different duplication mechanisms, distinct patterns emerged (**Supplementary Fig. S3** and **Table S6**). Tandem-duplicated gene pairs exhibited the highest proportion of conserved ACRs, the highest BLASTed ratios and the lowest mismatch rates (**Supplementary Fig. S3**), but this did not correspond to stronger expression correlation (e.g., dosage-balanced or paralogue dominance) (**Fig. 2B and Supplementary Fig. S1**). This discrepancy may reflect limitations of our current method, which tends to map ACRs back to themselves for tandem-duplicated genes (**Fig. 3A**), or it may result from other factors impacting transcriptional activities, such as positional effects, DNA folding or competition for transcription factors. In contrast, dispersed- and transposed-duplicated pairs displayed markedly different ACR genetic variation: low proportions of conserved ACRs, low BLASTed ratios, relatively high mismatch rates, and elevated fractions of unBLASTed ACRs (**Supplementary Fig. S3** and **Table S6**). Proximal-duplicated pairs, by comparison, exhibited variation patterns comparable to those of whole-genome duplication derived pairs, except with slightly higher BLASTed ratios (**Supplementary Fig. S3** and **Table S6**). These findings suggest that *cis*-regulatory elements are possibly more frequently co-duplicated with tandem and proximal duplicates, as they were with whole-genome duplicates in which *cis*-regulatory elements were entirely duplicated (Shoemaker et al. 2006), whereas variation in the presence or absence of duplicated *cis*-regulatory elements appears more common in transposed and disperse duplicates (Qiao et al. 2019). Collectively, these results indicate that different duplication mechanisms likely give rise to different types of *cis-* regulatory variants, thereby shaping transcriptional divergence differently.

### Transcriptional and ACR dynamics of duplicated gene pairs across tissues

By assigning each gene pair an expression pattern - non-expressed, “others”, mono-expressed, dosage balanced, paralogue dominance, specialization and divergence - across the seven examined tissues, we were able to characterize their cross-tissue transcriptional dynamics. In total, we identified 1,306 distinct expression combination patterns (**Supplementary Table S7)**. Among the top 86 most prevalent patterns, each represented by at least 100 gene pairs, the most common involved gene pairs that were mono-expressed in at least one tissue and non-expressed in the remaining tissues, collectively accounting for over 20% of all gene pairs (**Fig. 4A** and **Supplementary Table S7)**. Gene pairs with consistent mono-expression across all seven tissues also ranked high, representing the third most common pattern (**Fig. 4A** and **Supplementary Table S7**). Notably, in these pairs, the same gene was always consistently expressed across tissues (**Supplementary Fig. S4** and **Table S8**), aligning with previous observations that differential expression within gene pairs tend to occur in the same direction across tissues (Roulin et al. 2013). The second most common pattern was complete non-expression across all tissues, which is unsurprising given the limited number of tissues included in this study (**Fig. 4A** and **Supplementary Table S7**).

**Figure 4.**
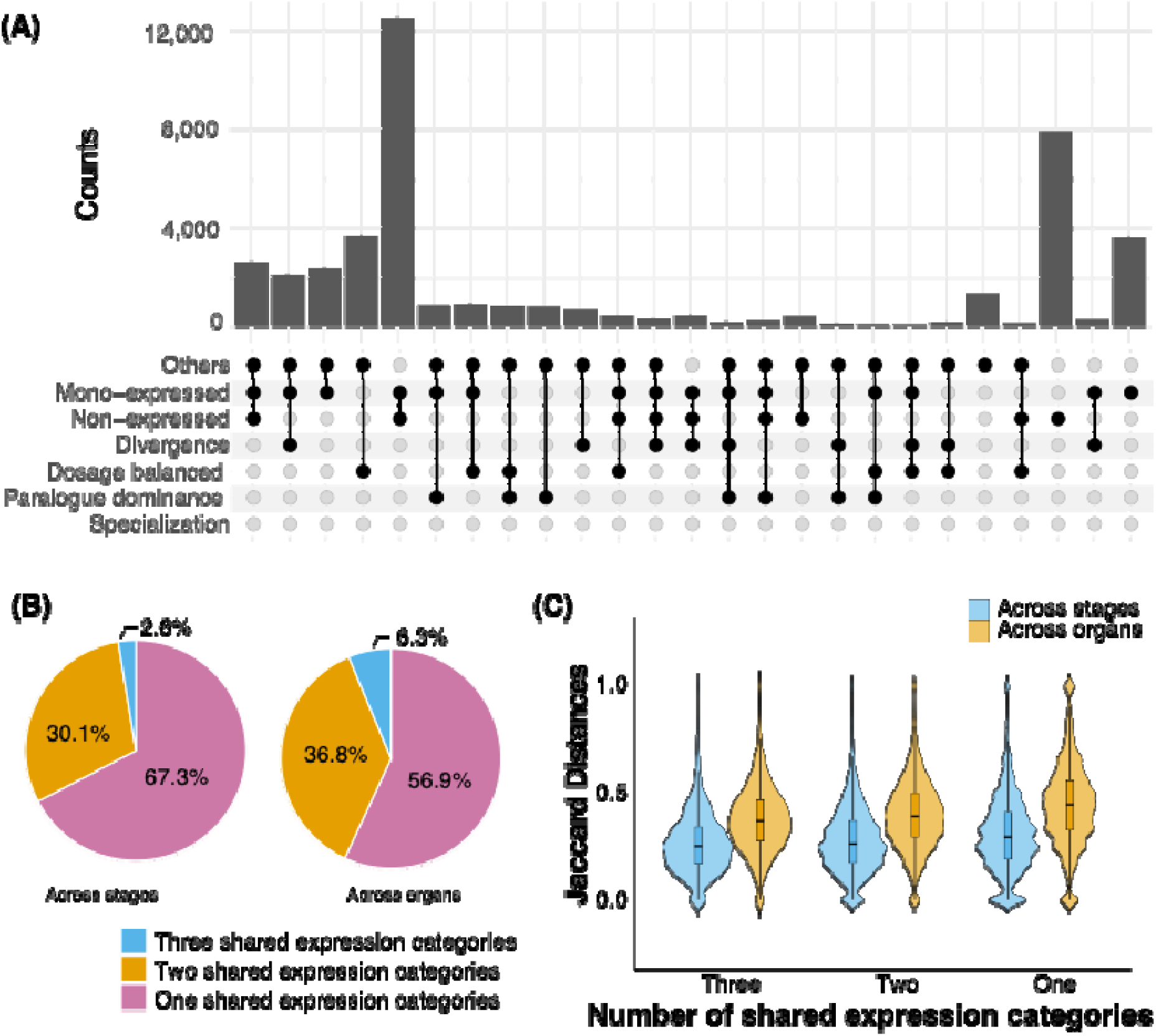
Across-tissue expression patterns of duplicated gene pairs. **A)** Number of duplicated gene pairs classified by combinations of expression patterns across seven examined tissues. **B)** Pie charts showing the percentages of duplicated gene pairs with different number of shared expression categories across stages or across organs. **C)** Jaccard distance, which measures the overlap of ACR chromatin accessibility profiles across stages or organs, of duplicated gene pairs group by the number of shared expression categories across stages or across organs.

The inclusion of multiple organs, root, hypocotyl, early nodule and seed, as well as seed sampled across distinct developmental stages, globular, heart, cotyledon, and early-maturation stages, further enabled us to assess the stability of gene pair expression patterns across both spatial and temporal dimensions. To facilitate interpretation, we consolidated expression patterns into four simplified categories: non-expressed, mono-expressed, correlated-expressed (dosage balanced, paralogue dominance, and specialization), and uncorrelated-expressed (divergence and “others”) (**Supplementary Table S9**). For each gene pair, we quantified the number of unique shared expression categories observed across organs or stages (**See Methods for details**). More than half of the gene pairs had a single shared expression category across stages (67.3%) and organs (56.9%) (**Fig. 4B**), indicating generally high consistency of expression patterns across tissues. Notably, 47% of gene pairs shared a single category across both stages and organs, although many of these were non- or mono-expressed (**Supplementary Table S9**). Fewer gene pairs shared a single category and more shared two or three categories across organs than across stages (**Fig. 4B**), suggesting greater transcriptional dynamics across organs. When examining expression stability of gene pairs grouped by their duplication mechanisms, these differences between across stages and across organs remained largely consistent; however, gene pairs derived from regional duplications exhibited a markedly higher proportion of only one shared category, compared with those derived from whole-genome or dispersed duplications (**Supplementary Table S10** and **Fig. S5**).

While gene pairs exhibit varied expression patterns across tissues, the genetic variants within their associated ACRs are determined by the underlying DNA sequence and remain unchanged across stages or organs. Consequently, these cross-tissue dynamics is unlikely to be driven directly by such genetic variants, aside from potential *trans-* effects, but rather by whether the same or different ACRs are accessible across tissues. We quantified this variation using the Jaccard distances, which measure the overlap of ACR accessibility profiles across stages or organs, with higher values indicating less shared accessibility (**Supplementary Tables S11-S12**). Surprisingly, the Jaccard distances showed little correlation with the number of shared expression categories across either stages or organs (**Fig. 4C**). Nevertheless, the higher Jaccard distances across organs than across stages (**Fig. 4C**), together with the greater expression dynamics observed across organs (**Fig. 4B**), suggests that variation in ACR accessibility profiles across tissues influences expression stability.

### Tracing the evolution of duplicated *cis*-regulatory regions following two rounds of whole-genome duplication

Soybean is a paleopolyploid species that has experienced two rounds of whole-genome duplication, theoretically giving rise to four duplicated genes sharing a single ancestral gene. Although extensive gene loss followed these duplications (Schmutz et al. 2010; Du et al. 2012; Zhao et al. 2017), the retained complete gene sets provide a valuable framework for investigating the evolution of *cis*-regulatory regions, duplicated alongside their associated genes (Li and Schmitz 2025), and its impact on gene expression. We focused on four-gene sets, each derived from a single ancestral gene shared across all bean-like (papilionoid) legume species, identifying a total of 1,904 gene sets, comprising 7,616 genes (**Supplementary Table S13**). By leveraging phylogenetic trees of individual gene families across papilionoid species, we classified gene pairs as originating from either the recent or ancestral whole-genome duplication (**Supplementary Table S13**). Recently duplicated gene pairs exhibited higher similarity in protein sequences and expression profiles, compared with ancestrally duplicated ones (**Supplementary Figs.6-7 and Table S14-S21**). As expected, ACR sequences associated with recently duplicated genes pairs were also notably more similar than those associated with ancestrally duplicated pairs (**Supplementary** Fig.8 **and Table S22-28**).

Four-gene sets exhibited complex expression patterns, reflecting both the number of expressed genes and the varied expression correlations among them. To characterize this expression divergence, gene sets were first categorized by the number of expressed genes – mono-, dual-, tri-, and quad-expressed, with dual expressed sets further subdivided into dual-recent (both expressed genes recently duplicated), and dual-ancestral (both ancestrally duplicated). Quad-expressed sets were the most frequent across most tissues, except in root, where non-expressed sets predominated (**Fig. 5A and Supplementary Table S29**). Dual-ancestral sets were consistently the least frequent, while other categories occurred at intermediate levels (**Fig. 5A and Supplementary Table S29**). Dual-recent sets always showed a strong skew toward high coexpression, reflecting their high expression similarity (**Supplementary Fig S9 and Table S30**). In contrast, other multi-expressed sets generally exhibited bimodal coexpression distributions, indicating either coordinated or divergent expression patterns, with the direction and magnitude of skew depending on both the number of expressed gene and the tissue context (**Supplementary Fig. S9 and Table S30**).

**Figure 5.**
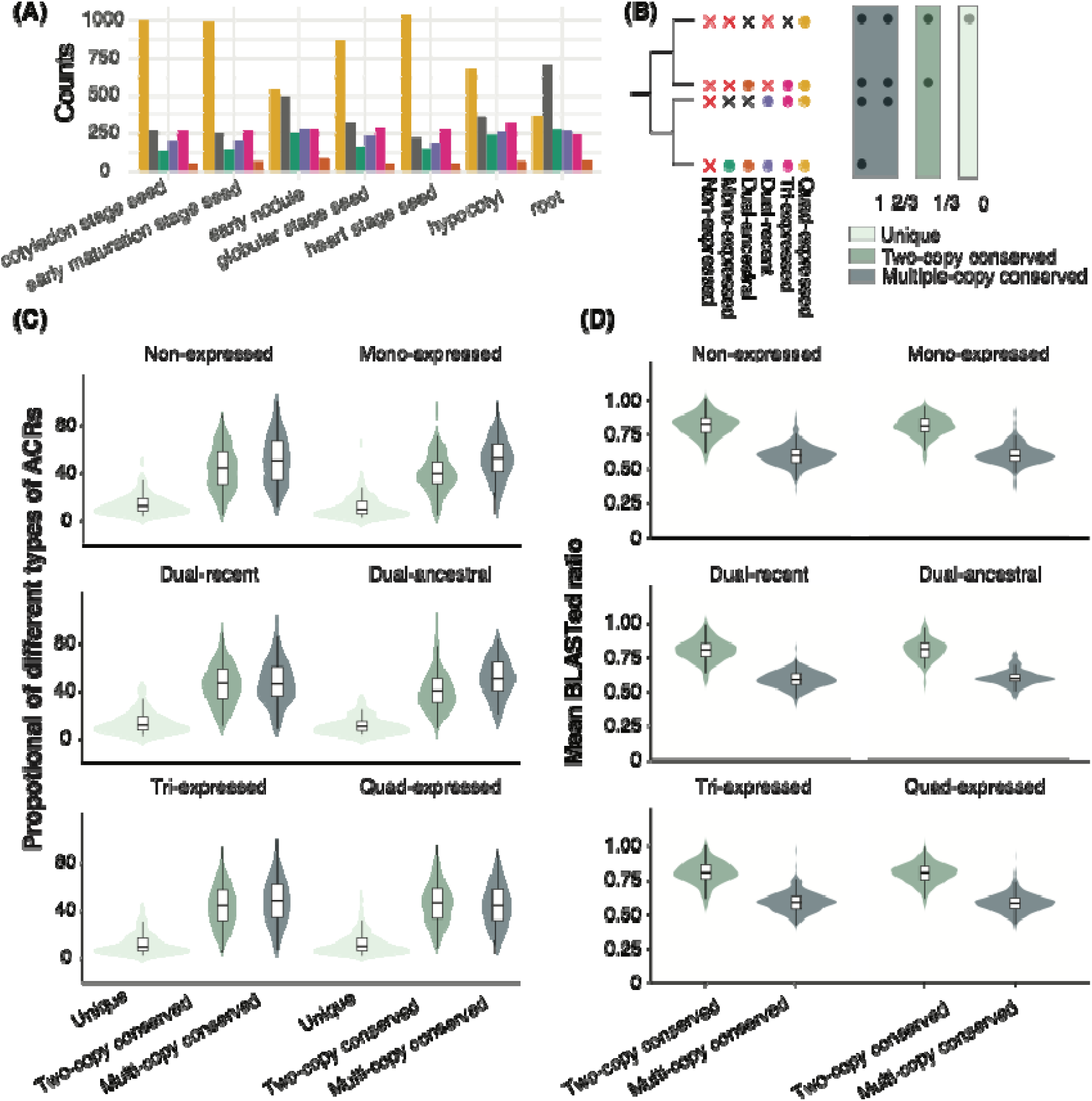
Proportions and sequence divergence of ACRs with different conservation states across gene sets with distinct expression patterns. **A)** Number of duplicated gene sets classified by the number of genes expressed, including, quad-expressed, tri-expressed, dual-recent, dual-ancestral, mono-expressed and non-expressed. **B)** Classifications of gene sets by the number of genes expressed and ACRs based on their conservation count: unique ACR, two-copy conserved ACR and multi-copy conserved ACR. **C)** Proportions of ACRs with different conservation states associated with gene sets exhibiting different expression patterns in cotyledon stage seed. **D)** Mean BLASTed ratio of two-copy conserved or multi-copy conserved ACRs across gene sets with different expression patterns in cotyledon stage seed.

To investigate the genetic variation and sequence origin of ACRs associated with these duplicated gene sets, we BLASTed each ACR associated with a duplicated gene against the reference regions of the other three duplicated genes within the same set. Each ACR was then assigned a conservation count, defined as the number of times it was conserved (BLASTed ratio > 0.1) across the three BLASTs. Based on this metric, ACRs were classified into three categories: unique ACRs, with no conserved sequences near any other duplicated genes, (conservation count = 0); two-copy conserved ACRs, conserved between only two genes, most often the two genes derived from the most recent duplication (conservation count = 1); and multiple-copy conserved ACRs, conserved across three or four duplicated genes after two rounds of duplication (conservation count = 3 or 2, with the latter suggesting possible loss of the ACR in one duplicate) (**Fig. 5B**).

The proportion of two-copy or multi-copy conserved ACRs was generally similar, and substantially higher than the proportion of unique ACRs, a pattern consistent across gene sets with varying expression profiles (**Fig. 5C, Supplementary Fig. S10 and Table S31**). These findings suggest that most ACRs associated with duplicated gene sets are conserved regions, likely retained from at least one round of whole-genome duplication, whereas only a small fraction are unique ACRs, potentially representing *de novo* regions that evolved at a low rate, although some unique ACRs may also result from loss in other duplicates. Notably, the relatively high frequence of multi-copy conserved ACRs was unexpected, given the extensive turnover of regulatory regions reported across multiple plant species with comparable divergence times (Lu et al. 2019). One possible explanation is that the divergence of non-coding sequences may be more pronounced across species than within a single-genome; however, this question requires further exploration in additional species and comparative contexts. We further calculated the average BLASTed ratio and mismatch rate for two-copy and multi-copy conserved ACRs. The higher genetic variation observed in multi-copy compared with two-copy conserved ACRs supports a temporal model of regulatory sequence divergence, in which ACRs gradually accumulate sequence variation over time (**Fig. 5C, Supplementary Fig. S11 and Table S31**). Overall, however, there were no substantial differences in either proportion or genetic variation of these ACRs among gene sets with different expression patterns (**Fig. 5B - 5C, Supplementary Fig. S11 and Table S31**).

### Evolution of cell-type-specific expression and cell-type-specific ACRs

Genes with cell-type-specific expression are essential for establishing or maintaining cell identity, with *cis*-regulatory elements determining the cell types, developmental stages and expression levels in which these genes are expressed (Wittkopp and Kalay 2012; Schmitz et al. 2022; Marand et al. 2023). We defined cell type-specific genes as those showing significantly higher expression in a given cell type compared to an equal number of randomly sampled cells from other cell types within the same tissue (Zhang et al. 2024). On average, 8,278 cell-type-specific genes were identified per tissue, with heart stage seed (11,497) having the highest number and early nodule (4,323) the lowest (**Supplementary Table S32**). Across developmental stages, these gene exhibited distinct dynamics: many maintained consistent cell-type-specific expression across stages (**Fig. 6A** - pattern 3), others were restricted to a single stage (**Fig. 6A** - pattern 2), and a smaller subset showed gains or losses of specificity during development (**Fig.** 6A - patterns 43 and 46), or shifts in specificity across stages (**Fig. 6A** - patterns 66 and pattern 102) (**Supplementary Tables S33-S34**).

**Figure 6.**
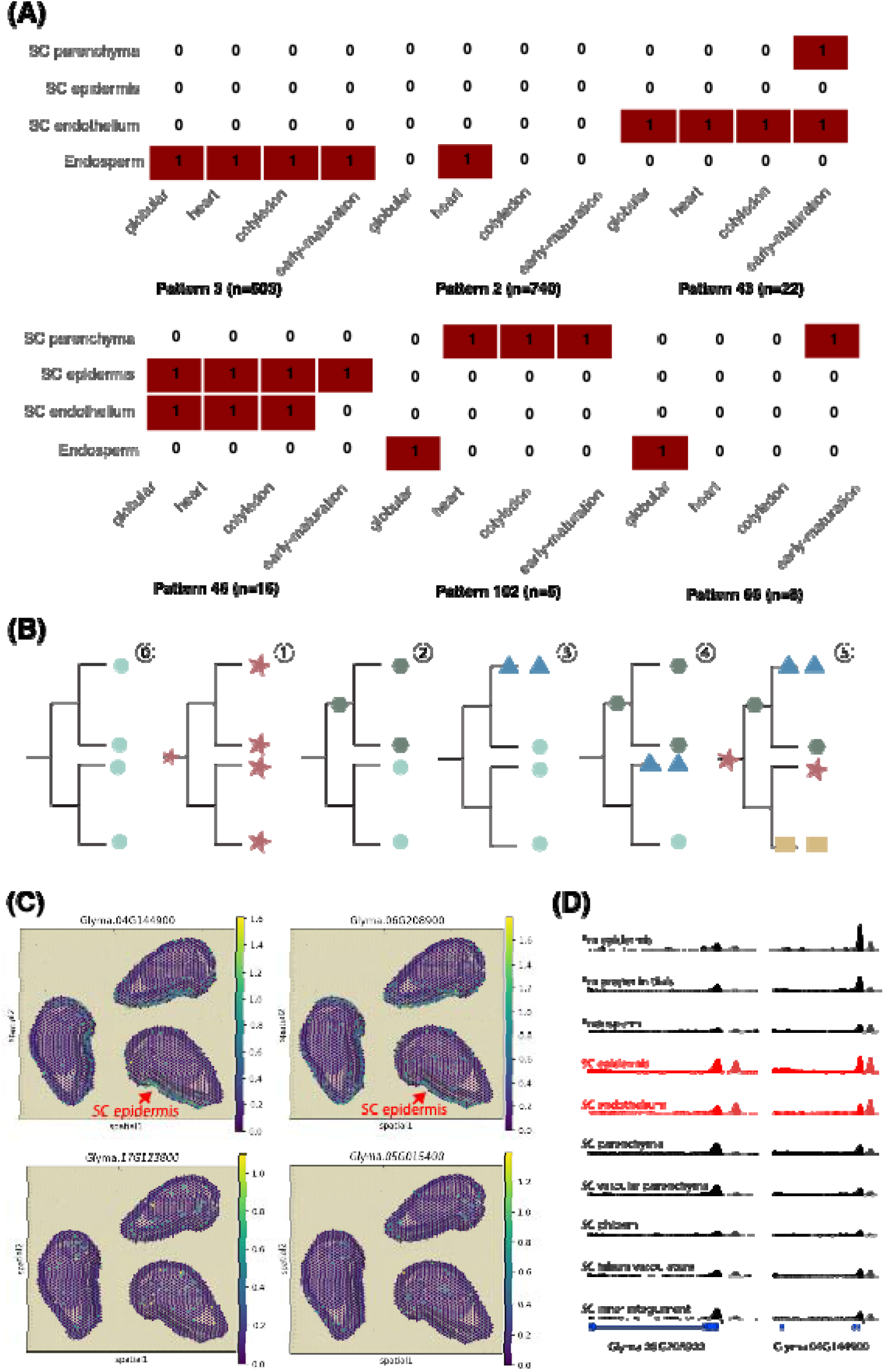
Evolution of cell-type-specific expression. **A)** Dynamics of cell-type-specific expression across stages, with ones indicating cell-type-specific expression in the given cell type and stage and zeros indicating absence of cell-type-specific expression. **B)** Schematic of initial expression l1 in duplicated gene sets and possible evolutionary trajectories of cell-type-specific expression: 11 evolved before the first round of whole-genome duplication; 11 evolved before the second round of whole genome duplication; 11 evolved after the second round of whole-genome duplication; 11 and 11 combinations of scenarios. **C)** Gene set in which two recently duplicated genes show seed coat epidermic-specific expression (top), while the other two do not (bottom). **D)** The ACRs associated with the duplicated genes exhibiting seed-coat epidermic specific expression showing higher chromatin accessibility in this cell type and seed-coat endothelium. Em: Embryo; SC: Seed Coat.

Placing these genes within the framework of four-gene sets - where duplicates initially shared the same gene body and regulatory regions - allows us to trace the evolutionary trajectories of cell-type-specific expression and potentially uncover the regulatory regions driving these changes. We focused on the four-gene sets in which all four genes were expressed and at least one exhibited cell-type-specific expression, thereby avoiding confounding effects from loss of expression in some duplicates. These duplicated genes were clustered using a binary matrix indicating the presence or absence of significant differential expression across cell types within the respective tissue (**see Methods**) (**Supplementary Figs. S12-S18**). Based on the number of shared clusters within each set and the ratio of genes assigned to each cluster, we inferred distinct evolutionary paths of cell-type-specific expression (**Fig. 6B**) (**Supplementary Table S35**). A few sets were assigned to only a single cluster, suggesting that cell-type specificity likely originated before the first round of whole-genome duplication and was subsequently maintained through both rounds of duplications (**Fig. 6B** 11) (**Supplementary Table S36**). In contrast, most sets had multiple clusters, reflecting the divergence or emergence of novel expression patterns, which may have arisen before (**Fig. 6B** 11) or after the second round of whole genome duplication (**Fig. 6B** 11); with some sets exhibiting combinations of these scenarios (**Fig. 6B** 11, 11) (**Supplementary Table S36**).

To explore the role of *cis*-regulatory activities in shaping these cell-type-specific expression dynamics within gene sets, we prioritized the analysis of their associated cell-type-specific ACRs (ctACRs) (Zhang et al., 2025). Specifically, we focused on ctACRs with significantly higher chromatin accessibility in the cell types where their associated duplicated genes were specifically expressed (**See Methods**) (**Supplementary Table S37**). Overall, conservation patterns of these ctACRs were broadly similar across gene sets with distinct evolutionary trajectories of cell-type-specific expression (**Supplementary Fig. S19**). Notably, gene sets forming a single cell-type-specific expression cluster tended to have a higher proportion of multi-copy conserved ctACRs compared to two-copy conserved or unique ctACRs (**Supplementary Fig. S19**). These findings suggest that ctACRs with different conservation states may play distinct roles in the evolution of cell-type-specific expression.

Importantly, by integrating cell-type-specificity with regulatory sequence conservation, we were able to pinpoint candidate ACRs responsible for cell-type-specific expression. For example, in one four-gene set, a recently duplicated pair displayed seed-coat epidermis specific expression in the cotyledon-stage seed (**Fig. 6C**, top panel), whereas the other pair lacked cell-type-specific expression (**Fig. 6C**, bottom panel). Within this set, we identified three ACRs exhibiting significantly higher chromatin accessibility in seed-coat epidermis cells (**Supplementary Table S37**). Two of them also showed elevated chromatin accessibility in seed-coat endothelium and were each associated with one of the cell-type-specific genes separately (**Fig. 6D**). Additionally, these two ACRs showed relatively high sequence conservation (**Supplementary Fig. S20**). Together, these results suggest that the regulatory elements within these two ctACRs likely arose *de novo* after the first round of whole-genome duplication, were duplicated during the second duplication, and were ultimately retained, contributing to cell-type-specific expression in these two genes (**Fig. 6C**. top panel). Although experimental validation is still needed to confirm their functions, this approach demonstrates a powerful strategy for identifying regulatory regions that drive cell-type-specific expression and for advancing our understanding of the evolution of cell-type specificity. Moreover, the dataset generated here provides a valuable community resource for investigating specific cell types or duplicated gene sets of interest.

## Discussion

### Single-cell genomics: a key to understanding transcriptional divergence and genetic variants in chromatin accessibility

The transcriptional divergence of duplicated genes has been extensively studied in polyploid plants (Du et al. 2012; Roulin et al. 2013; Zhao et al. 2017; Bird et al. 2020; Wang et al. 2021; Han et al. 2022; Fang et al. 2023; Hu et al. 2024). However, most previous studies relied on bulk RNA-seq, assessing divergence primarily through differential expression at the tissue level, with only few extending analyses to the single-cell scale (Coate et al. 2020). Compared to bulk approaches, single-cell genomics revealed more nuanced forms of divergence, including shifts in expression across cell types within a tissue and altered expression patterns across tissues (**Fig. 1 and 2**).

Characterizing both within-and across-tissue expression patterns provides a more comprehensive view of the spatiotemporal dynamics of gene expression. These multi-dimensional patterns also allows us to identify different drivers of expression divergence: genetic variation within ACRs primarily shapes within-tissue divergence, whereas variation in ACR chromatin accessibility profiles has a greater impact on cross-tissue divergence. These findings suggest that *cis*-regulatory elements probably act predominantly at the cellular level, and such patterns can best be captured through single-cell genomics.

Admittedly, bulk ATAC-seq, which profiles ACRs at the tissue level, remains powerful for detecting broad genome-wide patterns. In some cases, combining bulk ATAC-seq with snRNA-seq could identify some candidate regulatory regions underlying specific transcriptional patterns, for example, conserved ACRs with high BLASTed ratio and low mismatch rate associated with dosage-balanced gene pairs or unBLASTed ACRs linked to mono- or non-expressed gene pairs. However, scATAC-seq provides cellular context for ACRs, which is particularly valuable for dissecting mechanisms of cell-type-specific expression. When integrated with snRNA-seq, scATAC can help identify candidate ctACRs responsible for conserved or shifted cell-type-specific expression among duplicated genes, as exemplified in our duplicated gene sets (**Fig**. **6C** and **6D**). Nevertheless, pinpointing the causal regulatory elements within these ctACRs and understanding how these elements interact remains challenging.

### The evolution of *cis*-regulatory elements

The duplication of *cis*-regulatory elements represents a key source of novel regulatory sequences (McDonald and Reed 2024; Li and Schmitz 2025). Similar to protein-coding sequences of duplicated genes, which often experience reduced selective constraints (Kondrashov et al. 2002), duplicated *cis*-regulatory elements may also undergo relaxed selection, and accumulate mutations through genetic drift, potentially driving phenotypic novelty. This process is likely more pronounced following whole-genome duplications than small-scale duplications. Duplicates arising from whole-genome duplication tend to retain shared *cis-*regulatory elements (Shoemaker et al. 2006), whereas those from small-scale duplications may not (Arsovski et al. 2015). Consistent with this, we observed a higher percentage of ACRs that can be BLASTed to corresponding duplicated regions in whole-genome duplication derived gene pairs compared to small-scale duplication derived gene pairs, particularly transposed and dispersed duplicates (**Supplementary Fig. S4**).

On other hand, transposed or dispersed duplicated genes, where the gene body and likely its associated regulatory elements are translocated to new genomic environments, may come under the influence of regulatory elements pre-existed in their new surroundings. Alternatively, *de novo* regulatory elements could arise from sequences that were previously non-regulatory (Katikaneni and Lowe 2025; Li and Schmitz 2025). Both processes likely contribute to the greater divergence expression patterns observed in these duplicates. Together, modifications of duplicated elements and the *de novo* emergence of regulatory elements increase the diversity of regulatory sequences Moreover, our observation that most ACRs had its duplicated counterparts remained in the genome, with only a small subset of ACRs evolving *de novo*, suggest that retention and modification of existing elements might occur more frequently than *de novo* emergence.

Nevertheless, the precise genetic mechanisms and *cis*-regulatory activities that drive distinct expression outcomes – such as paralogue dominance versus specialization – remain unresolved. Still, other mechanisms, including epigenetic modifications such as DNA methylation and histone modifications, can also attribute to expression divergence. On the other hand, the relative contributions of evolutionary forces, including drift and selection, in shaping variation within existing *cis*-regulatory elements or in facilitating the emergence of novel ones are not yet fully understood. Finally, the interplay between the evolution of *cis*-regulatory elements and their associated protein-coding sequence remains underexplored.

### Cell-type-specific expression in the framework of duplicated genes

The four-gene sets involved in cell-type-specific expression used in this study provide a powerful framework for understanding the evolution of cell-type-specific expression. The diverse evolutionary trajectories of cell-type-specific expression were systematically explored and classified (**Fig.** 6B). Practically, this framework also enables the identification of candidate ACRs underlying cell-type-specific expression, as exemplified by seed-coat epidermal specific gene sets (**Fig. 6C** and **6D**). Identifying these regulatory sequences not only advances our understanding of cell identity formation, but also has potential application in synthetic biology, for instance, in designing regulatory regions to drive expression in specific cell types.

Furthermore, these dynamics of cell-type-specific expression provide a valuable perspective for studying subfunctionalization and neofunctionalization, the canonical outcomes of duplicated genes over long evolutionary time (Ohno 1970; Lynch and Conery 2000; Birchler and Yang 2022; Kuzmin et al. 2022; Gout et al. 2023). A persistent challenge has been defining or quantifying these processes, but changes in gene expression, particularly cell-type-specific changes, can serve as informative indicator. For example, Tran et al. (2025) demonstrated that reduced expression of one duplicated BLADE-ON-PETIOLE (BOP) gene in the bract-initiating region of *Capsella rubella* impaired its ability to regulate floral organ number and suppress bract formation, representing a step toward subfunctionalization. Building on this concept, our examination of cell-type-specific expression dynamics in duplicated genes provides a more quantitative means of evaluating these evolutionary outcomes. Integrating single-cell genomics with the comparative studies, essential for inferring ancestral expression patterns, offers a promising path to dissect subfunctionalization and neofunctionalization in greater detail.

In summary, understanding the evolution of paralogs is crucial, as many crops, including Solanum, grasses, and legumes, have similarly complex gene families that undoubtedly influence agriculturally important traits. Gaining insights into the transcriptional divergence of duplicated genes and the underlying *cis*-regulatory dynamics can inform strategies for fine-tuning gene dosage, facilitating the engineering of quantitative variation, and overcoming redundancy barriers in functional studies.

## Methods

### Duplicated gene pairs and gene sets in soybean

We identified duplicated gene pairs in soybean using the DupGen_finder pipeline (https://github.com/qiao-xin/DupGen_finder), with common bean (*Phaseolus vulgaris*) as the outgroup reference. We first performed all-against-all BLASTp within the soybean genome and all-to-all BLASTp between the soybean and *P.vulgaris* genomes, using the parameters “--sensitive --max-target-seqs 5 --evalue 1e-10”. The resulting BLASTed results were further filtered to retain only hits with sequence identity > 40% and coverage >70%. To retain redundancy across duplication modes, we used DupGen_finder.pl script, which classified duplicated gene pairs into five categories: whole-genome duplication, tandem duplication, proximal duplication, transposed duplication and dispersed duplication (**Fig. 1**). Gene pairs located in scaffolds were excluded from further analyses.

To identify gene sets consisting exclusively of four whole-genome duplication derived genes - presumably originating from two rounds of whole-genome duplications in soybean - we used the legume gene family dataset from SoyBase (https://www.soybase.org/), specifically the file “legume.fam3.VLMQ.sup1A_hsh.tsv”, which contains gene ID information for gene families shared across all papilionoid legume species and the corresponding phylogenetic trees (“legume.fam3.VLMQ.sup1B_trees”) built for each individual family. We first filtered the dataset to retain only gene families containing exactly four duplicated genes in soybean, and then further refined by cross-referencing with our previously identified whole-genome duplication derived gene pairs to ensure all four genes were products of whole-genome duplication. Finally, we used a customized script to identify the gene pairs clustered within the same clade of the gene tree, suggesting they likely arose from recent whole-genome duplication.

### Expression dynamics of gene pairs or gene sets

Raw gene counts from seven tissues - root, hypocotyl, nodule, globular stage seed, heart stage seed, cotyledon stage seed and early maturation stage seed - were obtained from snRNA-seq (Zhang et al. 2024). Data from each tissue was analyzed separately. Only genes with two replicates per cell type within the same tissue were retained for downstream analyses. For each tissue, we calculated the Spearman correlation between replicates of cell type and excluded cell types with low correlation (0.75 or blow). To ensure consistency between cell types identified in the snRNA-seq and scATAC-seq datasets, we focused only on cell types shared by both. Lowly expressed genes were filtered using “filterByExpr” function, and the remaining gene counts were normalized using “cpm” function in edgeR (Chen et al. 2025). Genes with zero expression across all cell types within a tissue were classified as “non-expressed” and excluded from further analysis. The remaining genes were used to construct coexpression networks (Benoit et al. 2025) to assess whether duplicated gene pairs exhibit similar expression profiles across cell types within a tissue.

To quantify expression relationships between duplicated genes within the same pair, we calculated the absolute mean and standard deviation (SD) of log2-transformed fold change across cell types within the same tissue (Benoit et al. 2025). Following (Benoit et al. 2025), we classified duplicated gene pairs into six categories based on coexpression and variation in expression levels, introducing “mono-expression” as an additional category:

1. Dosage-balanced: coexpression > 0.9, mean log_2_ fold change <1, SD log_2_ fold change <1
2. Paralogue dominance: coexpression > 0.9, mean log_2_ fold change >= 1, SD log_2_ fold change <1
3. Specialized: coexpression > 0.9, mean log_2_ fold change >= 1, SD log_2_ fold change >=1
4. Diverged: coexpression < 0.5, mean log_2_ fold change >= 1, SD log_2_ fold change >=1
5. Mono-expression: coexpression = NA.
6. Others: expressed gene pairs that did not fall into any of the above categories.

To investigate the dynamics of these expression patterns across tissues—particularly across developmental stages and different organs—we consolidated them into four simplified categories: non-expressed, mono-expressed, correlated-expressed (including dosage balance, paralogue dominance, and specialization), and uncorrelated-expressed (including divergence and “others”). We then assessed the number of shared expression categories across seeds collected at four developmental stages (globular, heart, cotyledon, and early maturation) and across four organs (hypocotyl, root, early nodule, and heart-stage seed). Non-expression was treated as a distinct case: it was excluded when evaluating shared expression categories if co-occurring with any expressed category and was only assigned when a gene pair was non-expressed across all organs or stages.

To assess transcriptional divergence of four-gene sets, we treated the four genes as six gene pairs: two representing recently duplicated pairs (each clustered into separate clades in the phylogenetic trees of the respective gene families) and four representing ancestrally duplicated pairs (**Supplementary Table S13**). For each gene pair, we calculated coexpression values, the absolute mean and standard deviation of log2-transformed fold change, following the same approach used above to examine expression divergence.

### Identification of cell-type-specific genes in each tissue in cultivated soybean

Cell-type-specific genes were identified using differential expression analysis for each cell type within its respective tissue (Zhang et al. 2025). Gene counts were aggregated across nuclei of the same cell type and compared to gene counts of an equal number of randomly sampled cells from other cell types within the same tissue, using edgeR (Chen et al. 2025). Genes with a log_2_ fold change > 1 and an adjusted p-value < 0.05 were classified as highly expressed in a given cell type. A gene was defined as cell-type-specific if it was classified as highly expressed in at least one cell type within its tissue. To further characterize their dynamics across tissues, we examined cell-type-specificity across the same cell types - endosperm, seed-coat endothelium, seed-coat epidermis and seed-coat parenchyma - in seeds sampled at four different developmental stages (globular, heart, cotyledon and early-maturation).

To investigate the evolution of cell-type-specific expression among gene sets, we first filtered for gene sets in which four duplicated genes were expressed and at least one duplicate exhibited cell-type-specific expression. For these genes, we constructed a binary matrix representing the presence (1) or absence (0) of significant differential expression across cell types within the respective tissue. Rows with no variation across cell types (i.e. all 0s) were removed prior to clustering. Pairwise distances between genes were calculated using the Jaccard distance, defined as 1 minus the ratio of shared presences to total presences and absences between two genes. Agglomerative hierarchical clustering was performed using the Ward.D2 linkage method. The optimal number of clusters was evaluated using the elbow method (within-cluster sum of squares) and silhouette analyses (average silhouette width) based on hierarchical clustering with Ward’D2 method. Genes were then assigned to clusters by cutting the dendrogram at the selected value of k. The resulting clusters observed within the gene set reflect the potentially varied evolutionary trajectories of cell-type-specific expression.

When all four genes grouped into a single cluster, cell-type-specific expression was inferred to have evolved prior to the first whole-genome duplication (**Fig. 6B** 11). A two-cluster distribution with a 2:2 ratio indicated evolution prior to the second whole-genome duplication (**Fig. 6B** 11), whereas a 1:3 ratio suggested evolution after the second whole-genome duplication (**Fig. 6B** 11). A three-cluster distribution with a 1:2:1 ratio was interpreted as a combination of these scenarios (**Fig. 6B** 11). Finally, when four clusters with a 1:1:1:1 ratio were observed, cell-type-specific expression was considered to reflect a more complex integration of the above evolutionary processes (**Fig. 6B** 11).

### Genetic variation in ACRs associated with gene pairs or gene sets

ACRs identified in soybean was obtained from (Zhang et al. 2024). The gene closest to each ACR was defined as the gene-associated ACR, and this association was determined using bedtools (Quinlan and Hall 2010). BLAST analysis was performed to examine the sequence divergence of ACRs associated with the two genes within the same duplication gene pair. The sequence of the ACR associated with one duplicated gene was used as the query, while the sequence of the two adjacent genes from the other duplicated gene served as the reference (**Fig. 3A**). These sequences were isolated using the “getfasta” function in bedtools (Quinlan and Hall 2010). Then sequence comparisons were performed using BLASTn with the following parameters “-task blastn-short -evalue 1e-3 -max_target_seqs 4-word_size 7 -gapopen 5 -gapextend 2 -penalty -1 -reward 1 -dust no -outfmt 6” following (Mendieta et al. 2024).

ACRs were classified as conserved if more than 10% (approximately 50 bp) were BLASTed to the reference region. The mean percentage of conserved ACRs were calculated for each gene pair by dividing the number of conserved ACRs by the total number of ACRs associated with that pair and then averaged across gene pairs with the same expression pattern. The BLASTed ratio was calculated by dividing the length of the BLASTed sequence in the reference region by the length of the query ACR. The mismatch rate was calculated as the number of mismatches between the ACR and the BLASTed sequence, divided by the length of the BLASTed sequence in the reference region. The percentage of unBLASTed ACRs was calculated by dividing the number of ACRs could not be BLASTed to the reference region by the total number of ACRs associated with the gene pair. The average BLASTed ratio, mismatch rate, and percentage of unBLASTed ACRs across all the ACRs associated with the gene pair were calculated to represent the level of sequence divergence of ACRs for each gene pair.

When measuring ACR sequence divergence across duplicated genes with the gene sets, we classified the ACRs based on conservation count, defined as the number of times an ACR was conserved (with a BLASTed ratio > 0.1) across the three BLASTs. Based on this metric, ACRs were categorized as: unique ACRs, two-copy conserved ACRs and multi-copy conserved ACRs (**Fig. 5B**). We examined the compositions of these ACRs for ACRs associated with each gene set. For each gene set, we calculated the average BLASTed ratio and mismatch rate for two-copy conserved ACRs, and multiple-copy conserved ACRs associated with each gene set. When comparing the ACR explaining cell-type-specific expression, we specifically focused on cell-type-specific ACRs exhibiting higher chromatin accessibility in the cell types where their associated genes were specifically expressed (Zhang et al. 2024).

Jaccard distance was calculated as 1 minus the ratio of shared presences to total presences among all ACRs associated with each gene set across developmental stages (globular, heart, cotyledon, and early-maturation stage seed) or organs (hypocotyl, root, early nodule, and heart-stage seed), providing a measure of the overlap in ACR accessibility profiles, where higher values indicate lower overlap (i.e., less shared accessibility).

All scripts for data processing and analysis are available at the following GitHub repository: https://github.com/Xianglichina/soybean_duplicated_genes_single_cell.git. During the preparation of this manuscript, we used ChatGPT to refine code and improve readability and language. All content was subsequently reviewed and edited by the authors, who take full responsibility for the final version.

## Acknowledgement

We sincerely appreciate the insightful feedback from Dr. Jeffrey Doyle from Cornell University, Dr. Scott Jackson from University of Georgia, Dr. Steven Cannon from USDA-ARS, and the members of the Schmitz lab on this manuscript, particularly Cullan Meyer, Kevin Sun, Dr. Hao Zhang, Dr. Shan Liang and Dr. Mark Minow. This research was supported by the United Soybean Board (<u>2432-201-0102</u>) and the National Science Foundation (<u>IOS-1856627</u>) to R.J.S.

## Author contributions

X.L., X.Z., and R.J.S. designed the study. X.L. and X.Z. analyzed data. X.L. prepared figures. X.L. and R.J.S. wrote the manuscript.

## Notes

### Competing Interest Statement

The authors have declared no competing interest.

### Summary of Updates

We re-identified duplicated genes using a more rigorous choice of outgroups and parameters and re-performed all subsequent analyses accordingly. We also expanded the section on expression patterns, examining them across developmental stages and organs as two distinct dimensions, temporal and spatial, to better characterize the expression dynamics of duplicated genes. In addition, we dissected the associated ACR dynamics across stages and organs to evaluate how changes in ACRs contribute to cross-tissue expression divergence. Finally, we systematically characterized the evolution of cell-type-specific expression within the framework of duplicated gene sets sharing a common ancestral gene. We also demonstrated how integrating cell-type specificity with conservation of regulatory sequences can identify candidate regulatory regions driving cell-type-specific expression, providing a valuable community resource for future studies on cell-type-specific gene regulation. Collectively, these modifications and the addition of these experiments have reinforced our original conclusions and, in many ways, strengthened them.

