## supplementary_figures for "From duplication to divergence: single-cell insights into transcriptional and *cis*-regulatory landscapes" (from another version of this preprint: version 2, 2025-05-08)

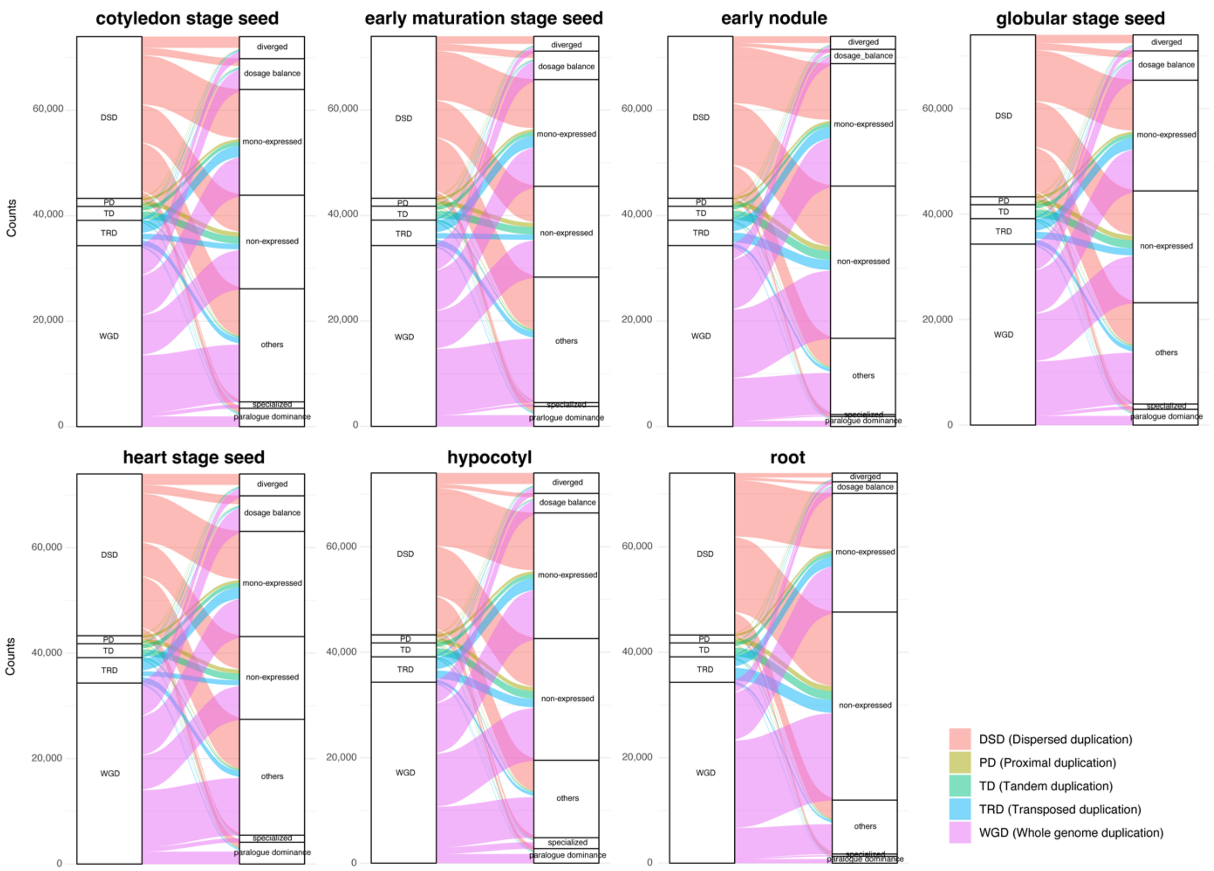


**supplementary fig. S1.** Relationship between duplication mechanisms and expression patterns within duplicated gene pairs across different tissues. The left panel categorizes genes based on duplication mechanisms: whole-genome duplication (WGD), tandem duplication (TD), proximal duplication (PD), dispersed duplication (DSD), and transposed duplication (TRD). The right panel represents gene expression patterns.

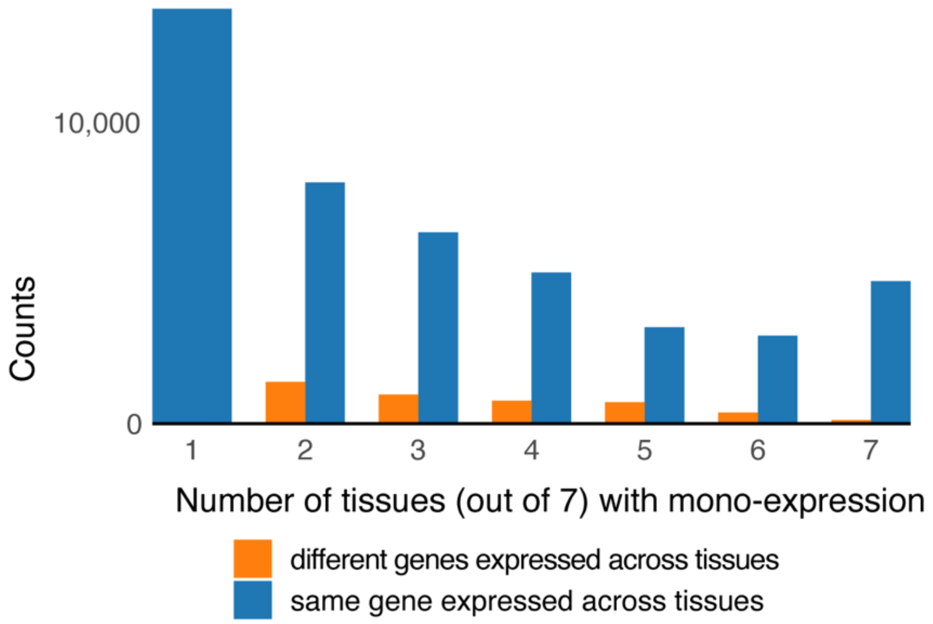


**supplementary fig. S2.** Mono-expressed gene pairs typically consistently exhibit the same gene being expressed while the other remain non-expressed across multiple tissues.

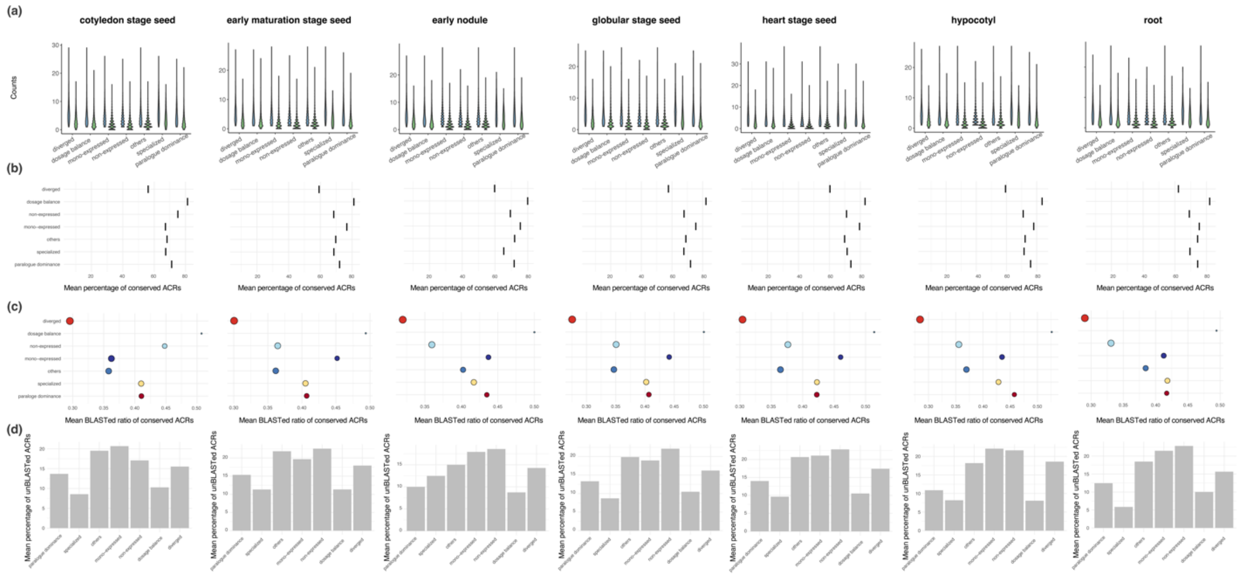


**supplementary fig. S3.** Genetic variations within ACRs associated with gene pairs with different expression patterns across different tissues. (a) Number of ACRs associated with each duplicated gene within the pairs exhibiting different expression patterns. Within each gene pair, the gene associated with a higher number of ACRs is classified into the group of large count while the gene associated with fewer ACRs is classified into the group of small count. (b) Mean percentage of conserved ACRs (black bars) across gene pairs with different expression patterns. (c) Mean BLASTed ratio of conserved ACRs across gene pairs with different expression patterns. The size of the dots represents the mean mismatch rate of conserved across gene pairs with different expression patterns. (d) Mean percentage of ACRs without a corresponding sequence in the reference region (unBLASTed ACRs) across gene pairs with different expression patterns.

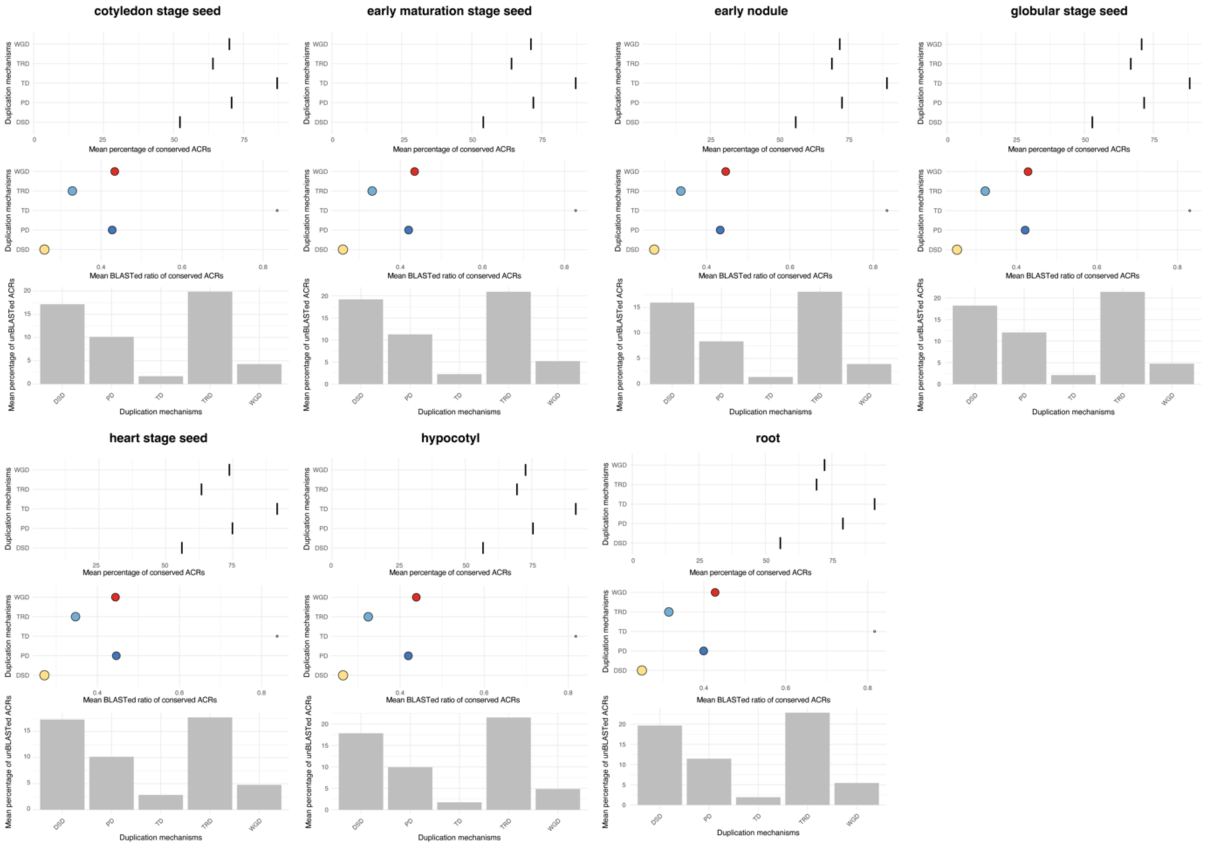


**supplementary fig. S4.** Genetic variations within ACRs associated with gene pairs derived from different duplication mechanisms. Top panel: mean percentage of conserved ACRs (black bars) across gene pairs derived from different duplication mechanisms. Middle panel: mean BLASTed ratio of conserved ACRs across gene pairs derived from different duplication mechanisms. The size of the dots represents the mean mismatch rate of conserved across gene pairs derived from different duplication mechanisms. Bottom panel: mean percentage of ACRs without a corresponding sequence in the reference region (unBLASTed ACRs) across gene pairs derived from different duplication mechanisms.

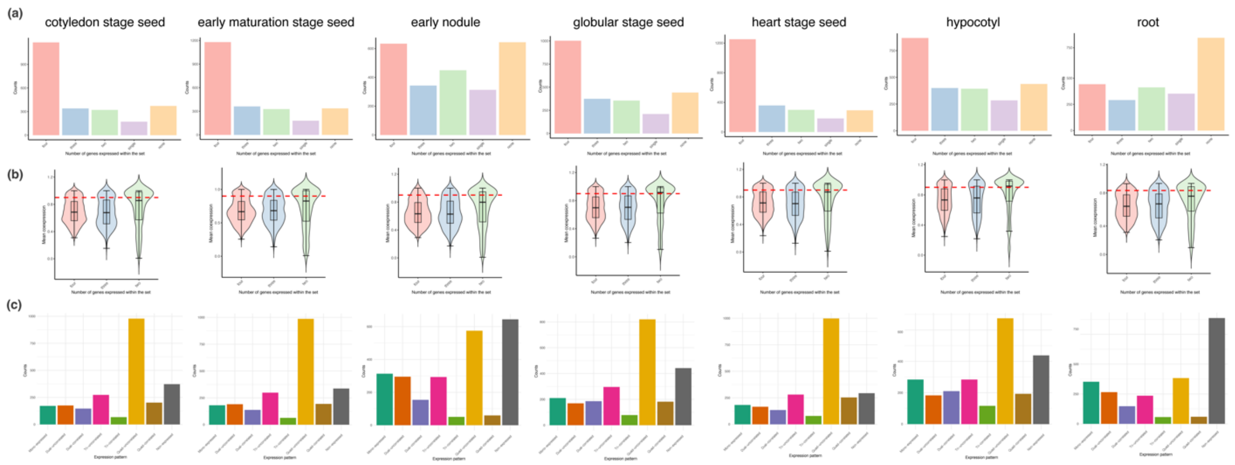


**supplementary fig. S5.** Transcriptional divergence across four whole-genome duplication derived duplicated genes sharing the same ancestral genes. (a) Counts of duplicated gene sets classified by the number of genes expressed with set in each tissue. (b) Mean cocorrelation among expressed genes within gene sets, grouped by the number of genes expressed per set in each tissue. (c) Counts of duplicated gene sets classified by expression patterns within each tissue, including mono-expressed, dual-uncorrelated, dual-correlated, tri-uncorrelated, tri-correlated, quad-uncorrelated, quad-correlated and non-expressed.

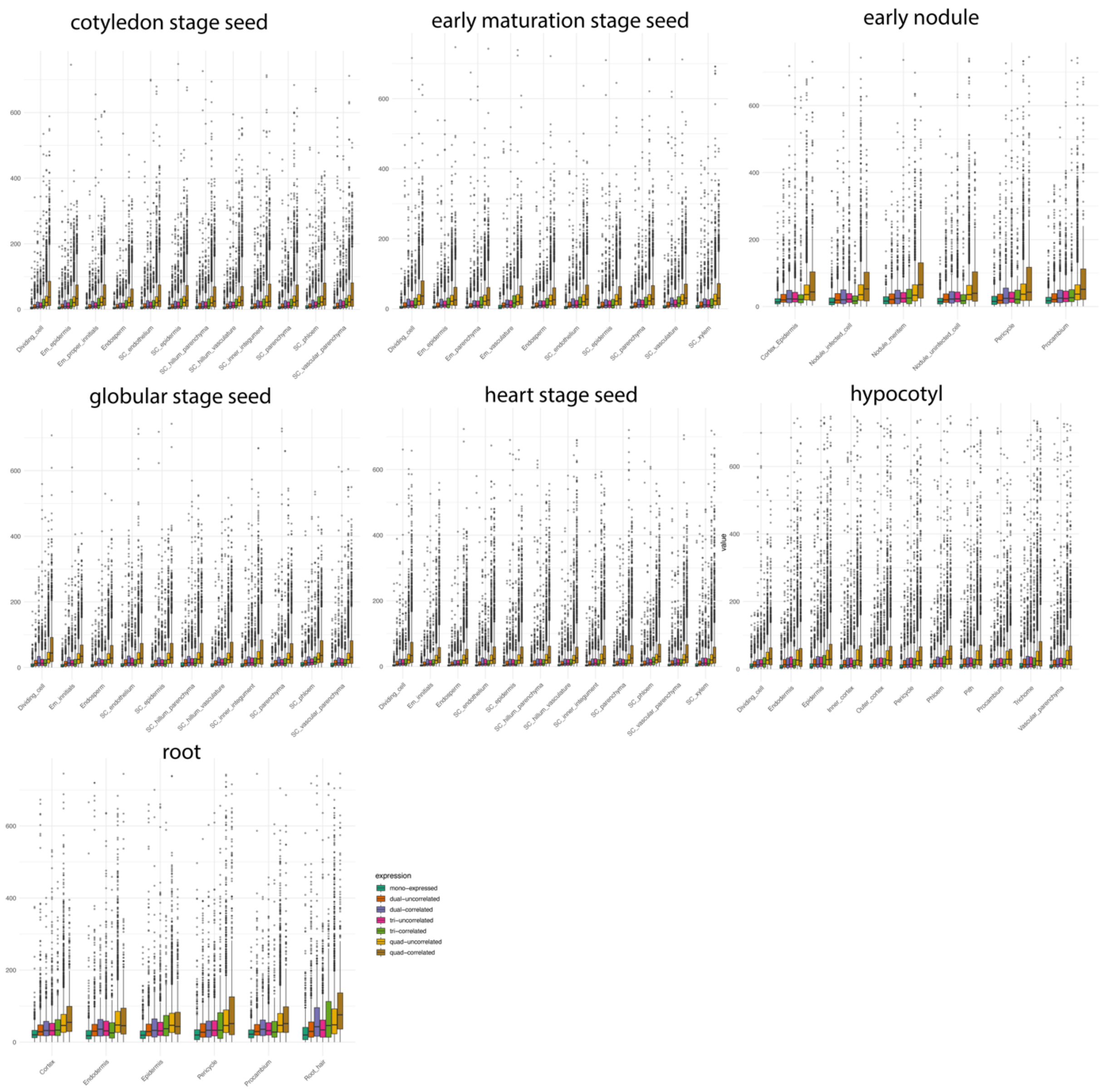


**supplementary fig. S6.** Expression levels of gene sets exhibiting different expression patterns across different tissues.

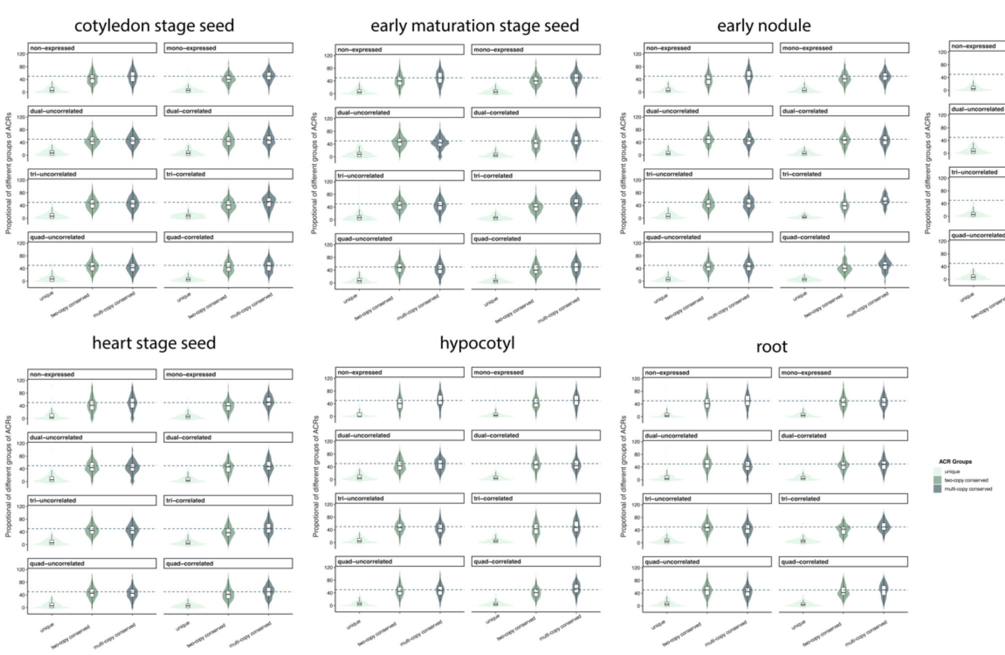


**supplementary fig. S7.** Proportions of each ACR category identified in gene sets with different expression patterns across different tissues.

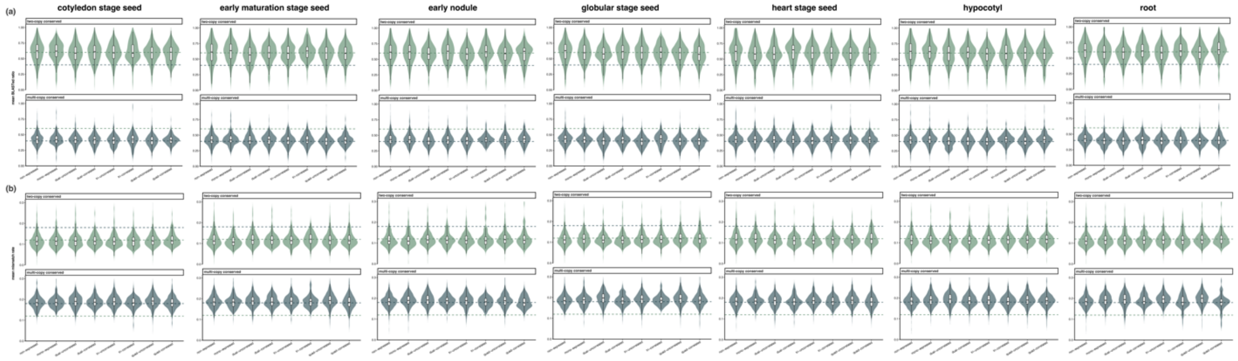


**supplementary fig. S8.** Measure of sequence similarity of ACRs associated with the duplicated genes within gene sets with distinct expression patterns across different tissues. (a) mean BLASTed ratio, and (b) mean mismatch rate.

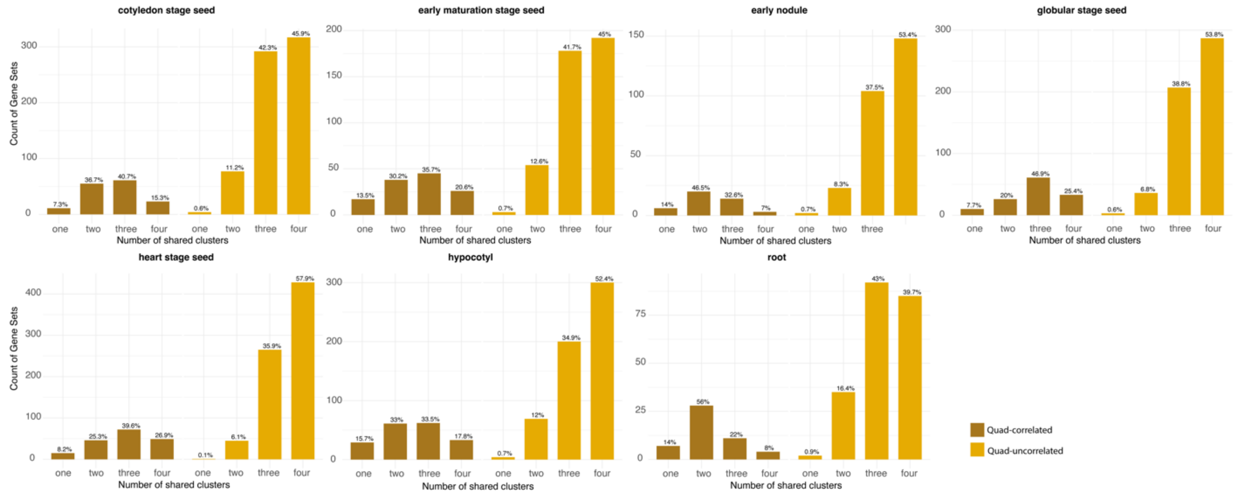


**supplementary fig. S9.** Counts of gene sets were classified based on the number of shared clusters identified within each set across different tissues. Gene sets exhibiting quad-correlated and quad-uncorrelated expression patterns were analyzed separately, and the percentage of each category was calculated relative to the total number of gene sets within each group.

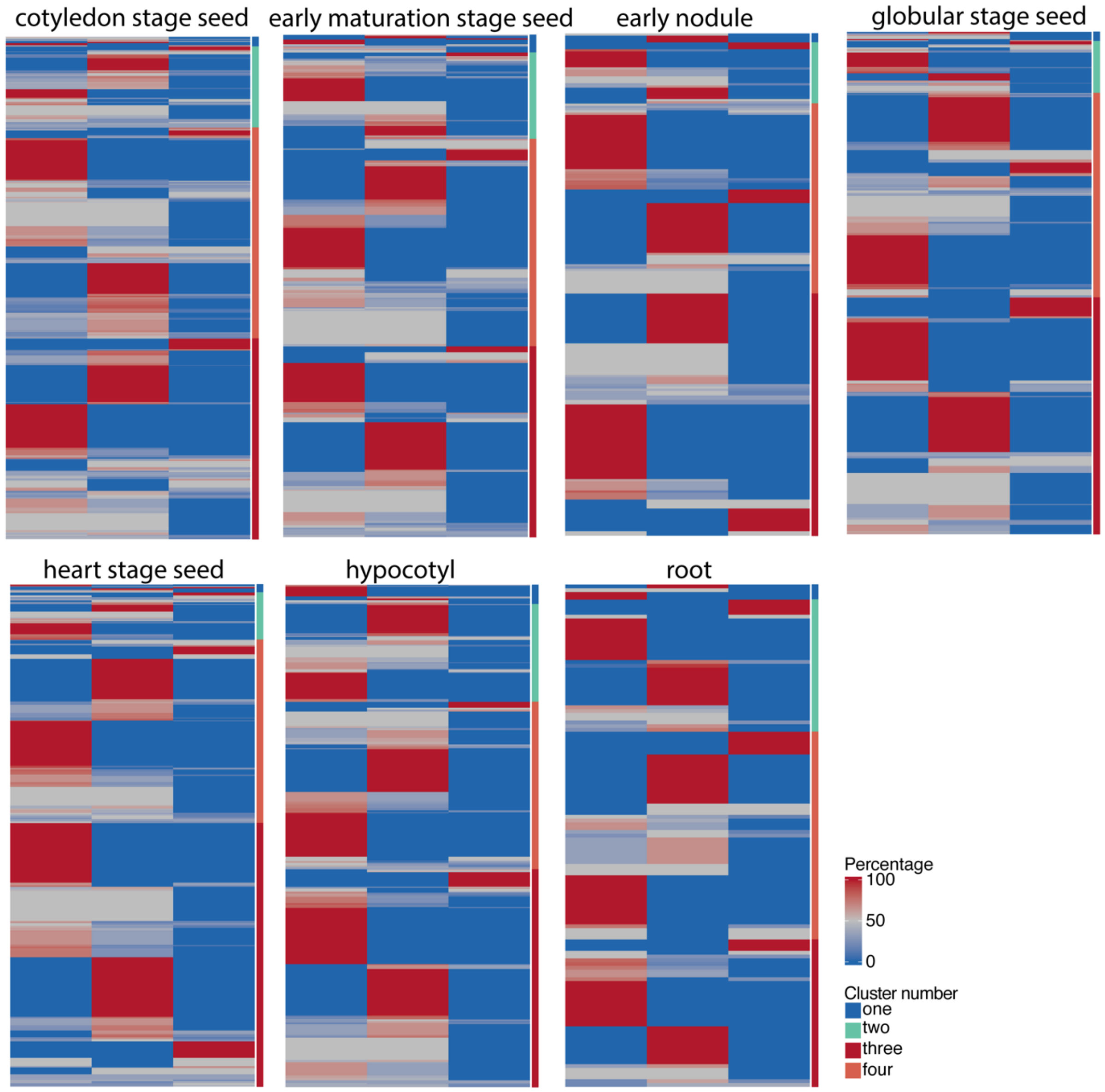


**supplementary fig. S10.** Heatmap of proportions of cell-type-specific ACRs (ctACRs) with different levels of conservation within gene set in different tissue types.

Each row represents a gene set in which all four genes are expressed and at least one exhibits cell-type-specific expression. The three columns display the proportions of two-copy conserved, multiple-copy conserved and unique ACRs associated with each gene set. Color intensity ranges from blue (0%) to red (100%). The bar alongside the heatmap indicates the number of shared clusters identified for each gene set.

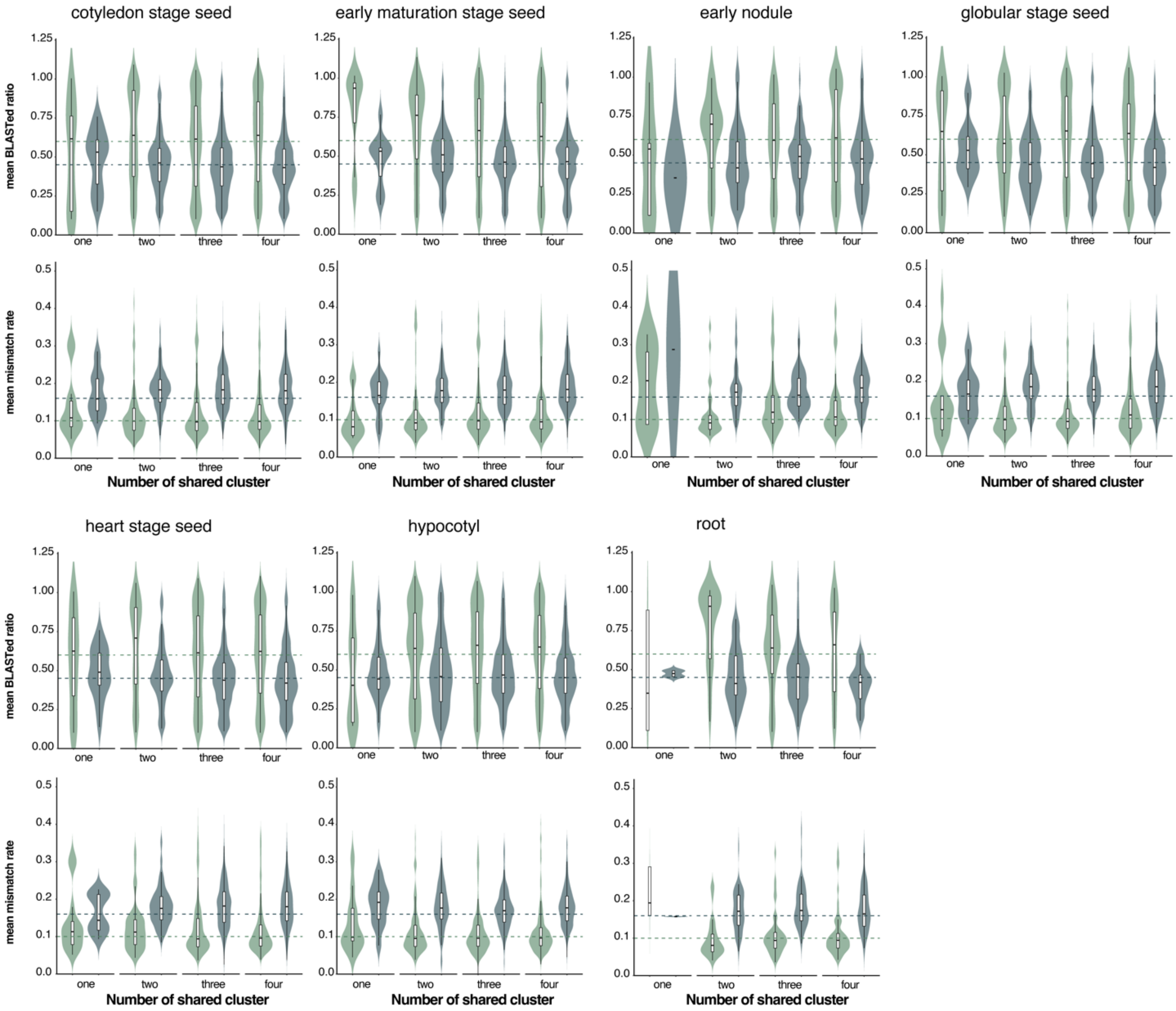


**supplementary fig. S11.** Measure of sequence similarity of ctACRs associated with the duplicated genes within gene sets with distinct number of shared clusters.

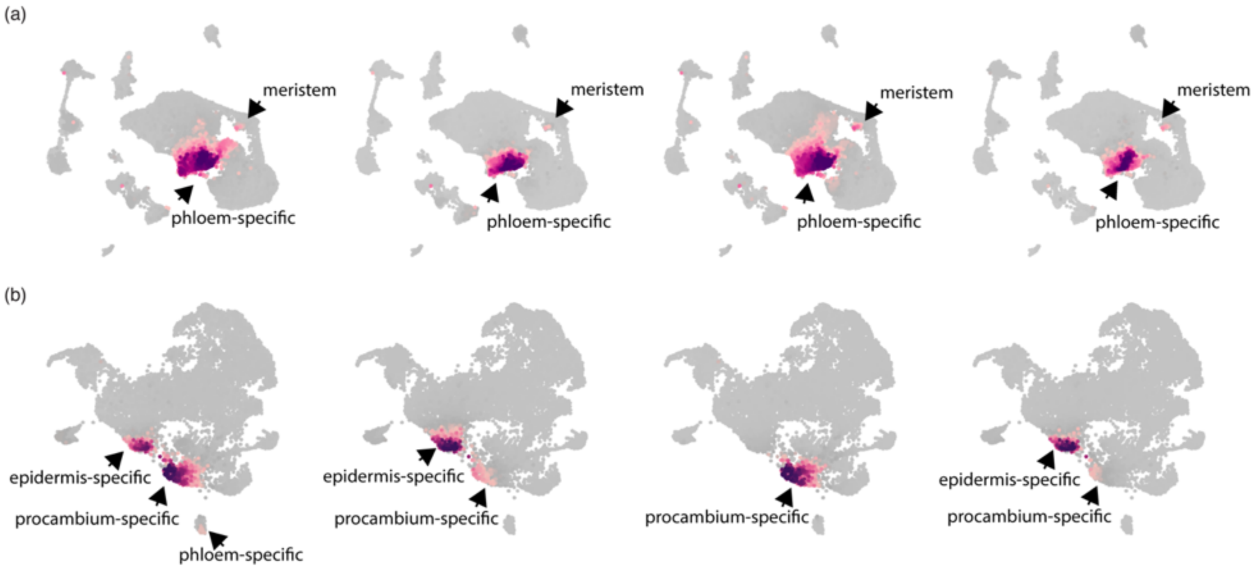


**supplementary fig. S12. A single gene set exhibiting distinct cell-type-specific dynamics in two different tissues.**

(a) Gene set exhibited phloem-specific expression in globular stage seed. (b) The same gene set exhibited varying cell-type-specific expression across different cell types in early-stage nodule tissue.
